# The value of behavioural activity records for conservation breeding: the case of Spix’s macaw in human care

**DOI:** 10.1101/2025.05.02.651864

**Authors:** Vladislav Marcuk, Cromwell Purchase, Katrin Scholtyssek, João Amador, Lorenzo Crosta, Petra Schnitzer, Heribert Hofer, Alexandre Courtiol

## Abstract

The Spix’s macaw (*Cyanopsitta spixii*), extinct in the wild since 2019, is currently maintained in *exsitu* breeding facilities. Captive breeding expanded the global population from 53 individuals in 2000 to ca. 400 by December 2025, enabling two soft releases. To support reintroduction, we introduce an ethogram of 85 behaviours and 1357 quantitative records based on observations of 123 birds at the ACTP Germany facility. Based on the analysis of the time-activity patterns of 10 pairs for 17 hours each, we also explore whether behavioural compatibility during the non-breeding season—measured here as time-activity similarity—relates to the breeding output of these pairs. Individual activity patterns were consistent, but females were markedly more synchronised with their mates than with other males. However, only pairs showing a high time-activity similarity bred successfully. Adjusting pair matching based on such similarity and improving general husbandry conditions (diet & enrichment) led to higher breeding output: pairs raising chicks increased from 0 in 2019 to 16 in 2024, fertility rates increased from 27% in 2019 to 64% in 2024, and annual chick output increased from 11 to 44 between 2019 and 2024. These results, based on 86 forced-mating attempts, support the idea that systematic behavioural monitoring can substantially contribute to enhancing conservation breeding in monogamous parrot species.

## 1. Introduction

The Spix’s macaw (*Cyanopsitta spixii*) is a monotypic psittacine of the Arini clade, currently classified as one of the five species that are extinct in the wild (Butchart et al., 2018; BirdLife International, 2026). After Johann Baptist von Spix first observed it about 200 years ago, Wagler formally described the species in 1832 (Barros *et al*., 2012). Historical sightings or reports remained rare for the past two centuries (Barros *et al*., 2012). In the 1980s, a tiny population of five individuals was rediscovered near Barra Grande and adjacent Riacho Melância in north Bahia (Roth, 1990a, 1990b). Between 1987 and 1988, the remaining three individuals from the only known population found their way into the illegal wildlife trade (Collar, 1992). A single male was discovered in the wild near Riacho Melância in July 1990, but no females were present. In 1995, a female held in captivity was released into the wild; she died shortly after, so this reintroduction attempt failed (Juniper, 2002). The last freeranging male was not seen after October 2000, leading to the classification of the species as extinct in the wild in 2019 (Butchart et al., 2018; BirdLife International, 2026) and to the initiation of a global *ex-situ* breeding program (Juniper, 2002). From an initial captive population of 53 individuals by December 2000 (Purchase, 2019), the population in human care grew to ca. 400 individuals as of December 2025. Despite this apparent success, little has been documented about the behaviour of the species and how the *ex-situ* breeding program and reintroduction attempts have likely benefited from such knowledge.

The first goal of this study was to provide a complete overview of behaviours in Spix’s macaw, including stereotypies and other behavioural disorders. Behavioural data are essential for establishing adequate husbandry guidelines (Luescher, 2006) and improving conservation breeding for reintroduction programs (Plair *et al.,* 2008; de Azevedo *et al.,* 2017). This includes eliminating stereotypies in birds in human care, as this is not only an issue of animal welfare but is also likely to affect the competence of individuals released into the wild. Much behavioural research has been dedicated to psittacines over the past decades, ranging from descriptions of behaviours (Dilger, 1960, Hardy, 1963, Buckley, 1968, Serpell, 1979; Levinson, 1980; Uribe, 1982; Lantermann, 1987; Rowley, 1990; Prestes, 1991; Schneider *et al*., 2006; Favoretto *et al*., 2024) to studies testing specific behavioural hypotheses or assessing complex behaviour paradigms in cognitive behaviour, communication or the effects of environmental enrichment (Pepperberg, 2000; Dahlin and Wright, 2007, Auersperg and von Bayern, 2019; Checon *et al*., 2020; Ramos *et al*., 2020). In macaws, detailed behavioural descriptions are only available for free-ranging populations for the Blue-winged macaw (*Primolius maracana*, Barros, 2001; in the wild) and the Red-fronted macaw (*Ara rubrogenys,* Christiansen and Pitter, 1992; Pitter and Christiansen 1995; 1997), and for populations in human care for the Scarlet macaw (*A. macao*, Uribe, 1982), Blue-and-gold macaw (*A. ararauna*; Uribe, 1982), and Lear’s and Hyacinth macaw (*Anodorhynchus leari* and *A. hyacinthinus*, Schneider *et al*., 2006; Favoretto *et al*., 2024). Hence, the behaviours of many New World parrots remain underexplored, limiting the scope for improvements in conservation breeding.

The second goal of this study was to establish a baseline of the time-activity patterns in Spix’s macaw under standard husbandry conditions. Time-activity patterns can differ substantially between conspecific individuals from *in-situ* and *ex-situ* populations (Cornejo *et al.,* 2005). Easy access to food and obvious differences in the local environment often induce shifts in time-activity patterns among parrots in human care. Previous studies indicate that captive birds spend their time predominantly resting or performing maintenance behaviours (Lantermann, 1998; Cornejo *et al.,* 2005; de Azevedo *et al*., 2016; Checon *et al*., 2020; Ramos *et al*., 2020). The increase in time spent resting at the expense of foraging, and an environment with reduced structure, are serious concerns, as they may be linked to the emergence of behavioural disorders such as stereotypies (Meehan *et al.,* 2004; Garner *et* al., 2006). Moreover, discrepancies in the time-activity patterns between captive and wild conditions are likely to impede reintroduction success. Released birds will indeed behave maladaptively if their behavioural plasticity is insufficient to readjust their activities to the demands of the wild environment (Cornejo *et al.,* 2005; Crates *et al.,* 2023). Establishing a baseline of time-activity patterns in Spix’s macaw will allow assessment of alternative enrichment protocols and help identify those that ensure that time-activity patterns of individuals in human care approach those of wild parrots.

Our third and final goal was to assess the relevance of behavioural compatibility for the *ex-situ* breeding output, which we did by examining the relationship between similarity in time-activity patterns between paired males and females and the breeding output of such pairs. Psittacine *ex-situ* programs often emphasise the importance of matching individuals to minimise inbreeding (Morrison *et al.,* 2020). However, pairings based solely on genetic criteria may be unsuccessful, as such pairing designs can trigger aggression and/or be associated with low breeding success in several psittacine species (Waugh and Romero, 2000; Luescher, 2006). Spix’s macaw is no exception, and once artificial insemination was discontinued in 2018, genetically informed pairings often resulted in infertile clutches or in individuals not showing any interest in breeding (see Results). Among the community of people breeding parrots, there is a consensus that behavioural compatibility is crucial for breeding success and that a high level of behavioural compatibility among partners is desirable. Unfortunately, there is very limited published information available on how behaviour relates to breeding success. If such a link existed, time-activity patterns might guide *ex-situ* breeding efforts and complement genetic choice criteria already in use.

## 2. Materials and Methods

### 2.1 Husbandry

To achieve our three goals, we recorded the behaviour of 123 Spix’s macaws on a weekly basis in the largest *exsitu* population in the world during 2018 and 2019. We also studied diurnal time-activity patterns and their similarity for 10 breeding pairs outside the breeding season (September-February) during those two years. The study took place in the facility of the Association for the Conservation of Threatened Parrots e. V. (ACTP) in Rüdersdorf, Brandenburg, Germany, where all Spix’s macaws are housed as pairs if adult or in flocks if the birds were still immature in partly isolated units. Each unit consists of 12 or 13 aviaries, which were subdivided into smaller subunits of 4–5 aviaries, each separated by a single indoor corridor. The subunits are insulated to reduce noise, so that only pairs located within the same subunit maintain auditory contact with each other while inside. Each aviary had an indoor and outdoor enclosure, with dimensions of 2 × 3.5 × 2.8 m and 16 × 2 × 3 m (length × width × height), respectively. The indoor aviary was heated to 18–21 °C from October to March and included a restricted selection of horizontal and diagonal perches (to encourage use of the maximum flight area), two feeding tables accessible from the corridor, an L-nest box, and various elements for environmental enrichment. The tiled floor in each interior enclosure was covered with a 2–3 cm thick layer of wood shavings. Each box was equipped with two high-definition cameras (Vicon V988D-W311MIR Dome Camera): one inside the unit and one inside the nest. These cameras recorded the birds’ activities for several consecutive weeks, with video files stored on a remote computer server. Outdoor enclosures consisted of individual constellations of perches and a canopy (1 m) that protected the birds from direct sun exposure and excessive rain. An artificial rain system was installed in all outdoor aviaries and operated on an automated schedule over the warmer months (April-September).

All birds were fed twice daily (08:00–09:00 AM and 15:30–16:30 PM) and supplied with additional pellets during the breeding season, which lasted from March–August. Food quantity was adjusted in the winter, and a maintenance diet for adults was implemented to counteract excessive weight gain and ensure the maintenance of birds close to desired body weights (female: 288 g, male: 318 g, average weights of *n* = 112 birds) during both semi-annual periods (breeding and nonbreeding). Water was provided *ad libitum*. At the beginning of the breeding season in March, the amount of food was increased, and vitamins and minerals were added. Further changes were implemented once pairs began to rear chicks.

### 2.2 Observation methods

Non-contact observations were conducted using video cameras to avoid behavioural changes caused by the observer’s presence. In total, 320 hours of video material were analysed to establish the behavioural repertoire, and 170 hours (17 hours per pair) were analysed to quantify time-activity patterns. The video sequences were stored externally (AVI format) and analysed with Avidemux (v. 2.7.4). Behaviours for the ethogram were categorised in ten broad categories (Fig. 1), which are described in detail in the Appendix I, including *maintenance behaviours* (all behaviours included in Fig. A2a, behaviour 1 to 12), *physiological behaviours* (see Fig. A2b, behaviour 1 to 5), *locomotion behaviours* (active forms; Fig. A2c, behaviour 1 to 3), *inactivity behaviours* (Fig. A2d, behaviour 1 to 3), *agonistic behaviours* (Fig. A2e, behaviour 1 to 13), *displacement behaviours* (Fig. A2f: behaviour 1 to 11), *submission behaviours* (Fig.. A2g: behaviour 1 to 11), *social behaviours* (Fig. A2h: behaviour 1 to 7), *sexual behaviours* (Fig A2i: behaviour 1 to 8) and *behavioural disorders* (Fig. A2j: behaviour 1 to 12). This classification expands a preliminary survey which focused on agonistic and submission behaviours (Marcuk *et al*., 2020). The behaviours of each individual were analysed over the full diurnal period (from 05:00 to 21:00, i.e., the light hours).

**Figure 1.**
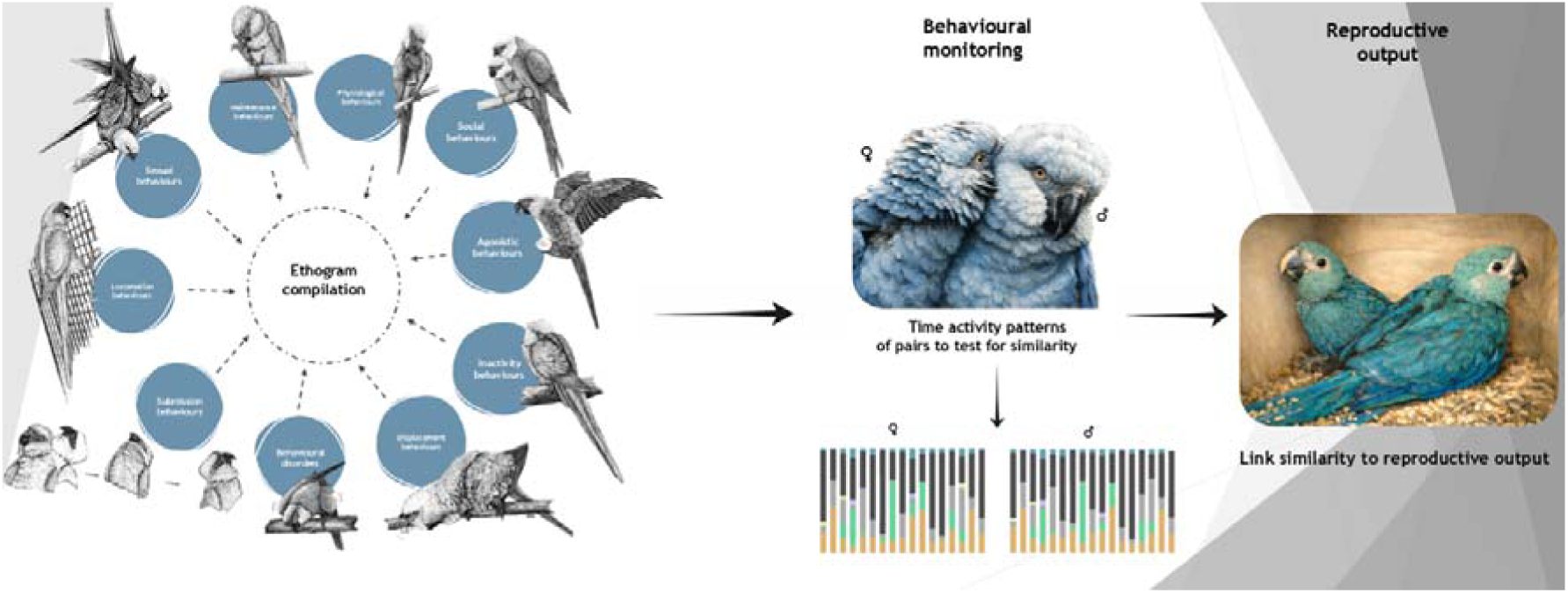
– Graphical abstract with the associated behaviour categories featuring each characteristic behaviour of the respective category. The time-activity patterns were compiled using some of the behavioural categories mentioned (illustrations by V. M.). Illustrations for agonistic and submission behaviour adapted from Marcuk *et al*., 2020. Illustration of a pair based on a picture from K.S.

**Figure 2.**
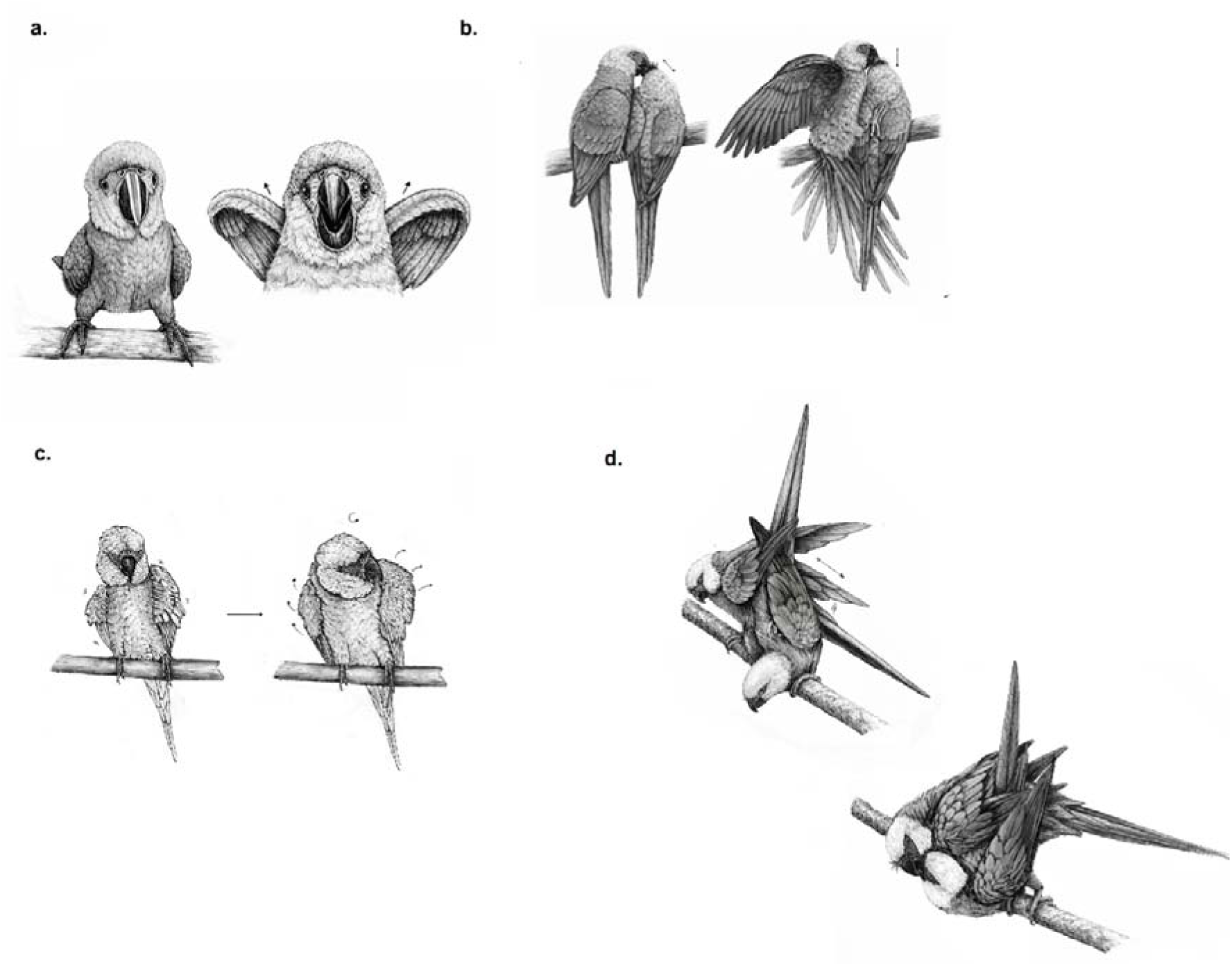
– Sample of illustrations of different behaviours: a.) begging, b.) sexual allo-feeding, c.) body shake, and d.) active phase of a copulation. The complete ethogram is provided in SI.

Time-activity patterns were recorded for ten pairs, where the duration of each behaviour was analysed for each hour, rounded to the nearest second and assigned to the respective behaviour category. Displacement behaviours were lumped together with submission or agonistic behaviours due to their very short duration. Since time-activity patterns were observed exclusively in indoor enclosures, neither heat display nor ruffling was observed. Therefore, physiological behaviours accounted for most of the active “foraging time” (e.g., food and water intake, defecation). Time activity patterns were also recorded during the non-breeding season. Therefore, behaviours related to sexual activity (social behaviours 4 to 7, sexual behaviours 1 to 8, and behavioural disorders 11 to 12) were excluded from the analysis of time-activity patterns.

We selected these pairs so as to capture a wide range of demographic history (e.g., aged between 4 and 22 years, individuals from different genetic lines = five of six founders represented, generations and birth-locations, and pairs that bred or didn’t breed), with the constraint that we could only retain pairs for which the male and the female were morphologically sufficiently different to unambiguously assign records to individuals (based on plumage aberrations, bare parts resulting from plucking, different iris colouration or, on some occasions, the colour and variation of the leg bands). All 20 individuals were hand-reared.

The interior was standardised across all indoor enclosures to minimise the impact of environmental factors on the birds’ behavioural repertoire or activity period during the observation period. None of the nest boxes were open, ensuring that none of the pairs in this study engaged in sexual behaviours. The observations were conducted during the early non-breeding season at the beginning of September in 2018 (three pairs; 5–8 September) and 2019 (seven pairs; 2–4 September). All individuals were adults (minimum age was 4 yrs). The observations took place before an enrichment plan was initiated in mid-September 2019. The recorded time-activity patterns thus constitute a baseline treatment without the influence of any sort of environmental enrichment. Breeding data were collected for the ten pairs between 2014 and 2024 at Al Wabra Wildlife Preservation and ACTP Germany. We recorded the number of eggs laid, the number of fertile eggs, and the number of weaned chicks produced for each female. Eggs and chicks resulting from artificial inseminations were not included. All eggs were inspected with a light source (i.e., candled) at least once to determine whether an embryo was developing or not. We also counted the total number of offspring once they were weaned.

### 2.3 Data analysis and statistics

To assess behavioural similarity, we excluded social behaviours, since interactions must necessarily occur in synchrony. We also excluded behavioural disorders since mates were never observed to mirror such behaviours. As mentioned above, both the timing of the experiment and its design precluded the expression of sexual behaviours. Using this data, we first performed a hierarchical cluster analysis using Ward**’**s method implemented in OriginLab 2025 (OriginLab Corporation, Northampton, MA 01060, US) to determine whether the actual pairs were identified as clusters based on their time-activity patterns. For this analysis, we used the following options: datatype = standardized data; type = Ward; average type = Euclidean; k = 10 clusters, equal to the number of pairs. We replicated these results using the R package pvclust (v. 2.2.0, Suzuki and Shimodaira, 2006) to assess dendrogram stability via non-parametric bootstrapping (1000 bootstrap replicates). Similarly, we used the R package vegan (v. 2.7.5, Okansen *et al.,* 2026) to test whether the 10 groups defined by the cluster analysis differed more behaviourally from one another than would be expected by chance, using the function ANOSIM with its default settings.

Second, we computed a similarity index for each pair to quantify the observed similarity between time-activity patterns. The similarity index *S* was defined as the sum of absolute differences between the percentages *p at* which each behaviour was displayed in each hour (hour_1_…hour_17_) of the male () and the female (), as:

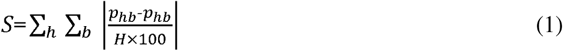

where *h* is the index of each hourly period between 05:00:00 and 21:59:59 (i.e., 05:00:00–05:59:59, 06:00:00– 06:59:59, …, 21:00:00–21:59:59), *b* is the index of each behaviour category considered (i.e., maintenance, foraging, submission, agonistic, resting & locomotion), and *H* is the total number of hourly periods recorded (here 17). A similarity index of 0 represents the lowest conceivable similarity of the time-activity behaviour patterns between female and male, while a value of 1 would represent pairs with identical time allocation across all six behaviours for every recorded hour. A high similarity in time-activity patterns therefore likely reflects a high behavioural synchrony between both partners.

We also computed *S* between each female and all ten males to examine how the similarity index of a female and its mate compared with the index between her and any other male. We compared the *S* values of females with their actual mates to the *S* values of females with all 10 males using an exact binomial test. Here, the null hypothesis was that a female’s actual mate was as likely as any other male to be the one showing the highest behavioural similarity to the female.

Descriptive statistics are presented as mean ± standard deviation (SD), with the range in parentheses. All statistical tests were performed in R (v. 4.3.4, R Core Team, 2024) with a two-tailed significance level of *a* = 0.05. To compare the overall percentages in time-activity patterns between the sexes, we used the exact Mann-Whitney U test as implemented in the R package coin (Hothorn *et al*., 2008).

## 3. Results

### 3.1 Ethogram

For the ethogram, we described a total of 85 behaviours categorised into 10 broad behaviour categories. In total, 1357 quantitative records were obtained. The ethogram included the following behaviour categories: *maintenance behaviours* (1. Body shake, 2. Head scratch, 3. Head shake, 4. Tail wag, 5. Wing & leg stretch, 6. Bilateral wing stretch, 7. Yawn, 8. Bill grind, 9. Bill wipe, 10. Touch-foot, 11. Autopreening, 12. Bath), *physiological behaviours* (1. Ruffling, 2. Heat-exposure display, 3. Drink, 4. Food intake, 5. Defecation), *locomotion behaviours* (1. Move, 2. Climb 3. Flying), *inactivity behaviours* (1. Perch, 2. Resting, 3. Roosting), *agonistic behaviours* (1. Neck & head feather raise, 2. Foot-lift, 3. Bill gape, 4. Wing-raise display, 5. Lunge, 6. Bite, 7. Bill fence, 8. Claw, 9. Rush, 10. Flying approach, 11. Flight attack, 12. Fight, 13. Redirected aggression), *displacement behaviours* (1. Displacement preen, 2. Displacement food-intake, 3. Displacement rub, 4. Displacement scratch, 5. Displacement hold-bite, 6. Displacement head down shake, 7. Displacement yawn, 8. Displacement allopreening, 9. Displacement mutual feed, 10. Irritated body shake, 11. Bill clasp), *submission behaviours* (1. Turn away, 2. Slide away, 3. Alert and fear reaction, 4. Apparent death display, 5. Bob, 6. Head-tilt solidarity display, 7. Crouch-quiver solidarity display, 8. Upside-down lift solidarity display, 9. Peer, 10. Unison jerk, 11. Singleton jerk), *social behaviours* (1. Contact-sitting, 2. Mutual nibbling, 3. Allopreening, 4. Reciprocal cloacal preen, 5. Non-sexual social play, 6. Begging, 7. Non-sexual Allofeeding), *sexual behaviours* (1. Sexual social play 2. Apparent copulations, 3. Jerk display, 4. Preen display, 5. Scratch display, 6. Rapid turn, 7. Sexual allofeeding 8. Copulation), *behavioural disorders* (1. Erratic flights, 2. Head tilt, 3. Crouch-quiver solidarity display, 4. Upside-down lift solidarity display, 5. Loop-walking, 6. Pterotillomania or feather plucking, 7. Overt allo-preening, 8. Auto-mutilation, 9. Allo-mutilation, 10. Redirected aggression, 11. Egg destruction, 12. Infanticide). Fig. 1 & 2 provides some illustrations. Detailed descriptions and methodological details are provided in the Appendix (AI).

### 3.2. Time-activity patterns

Table 1 summarises the proportion of time spent across all eight main activity categories during the observation period. In all monitored individuals, the predominant activity pattern was inactivity at 49.0 ± 4.9% (40.7–57.9%, Fig. 3), followed by maintenance at 18.1 ± 3.0% (10.5–23.6%), and social behaviour at 14.2 ± 2.1% (9.3– 16.3%). Physiological behaviours accounted for an average of 8.3 ± 1.8% (5.7–11.5%), and locomotion contributed on average 5.9 ± 1.6% (4.1–8.9%). Intra-pair aggression was documented only in a single pair, formed by individuals #59 and #91, during the observation period, but even for this pair, such behaviour remained rare and was the least common of all categorised social behaviours. No significant sex-specific differences were observed for any of the enlisted behaviour categories (see Table 1).

**Figure 3.**
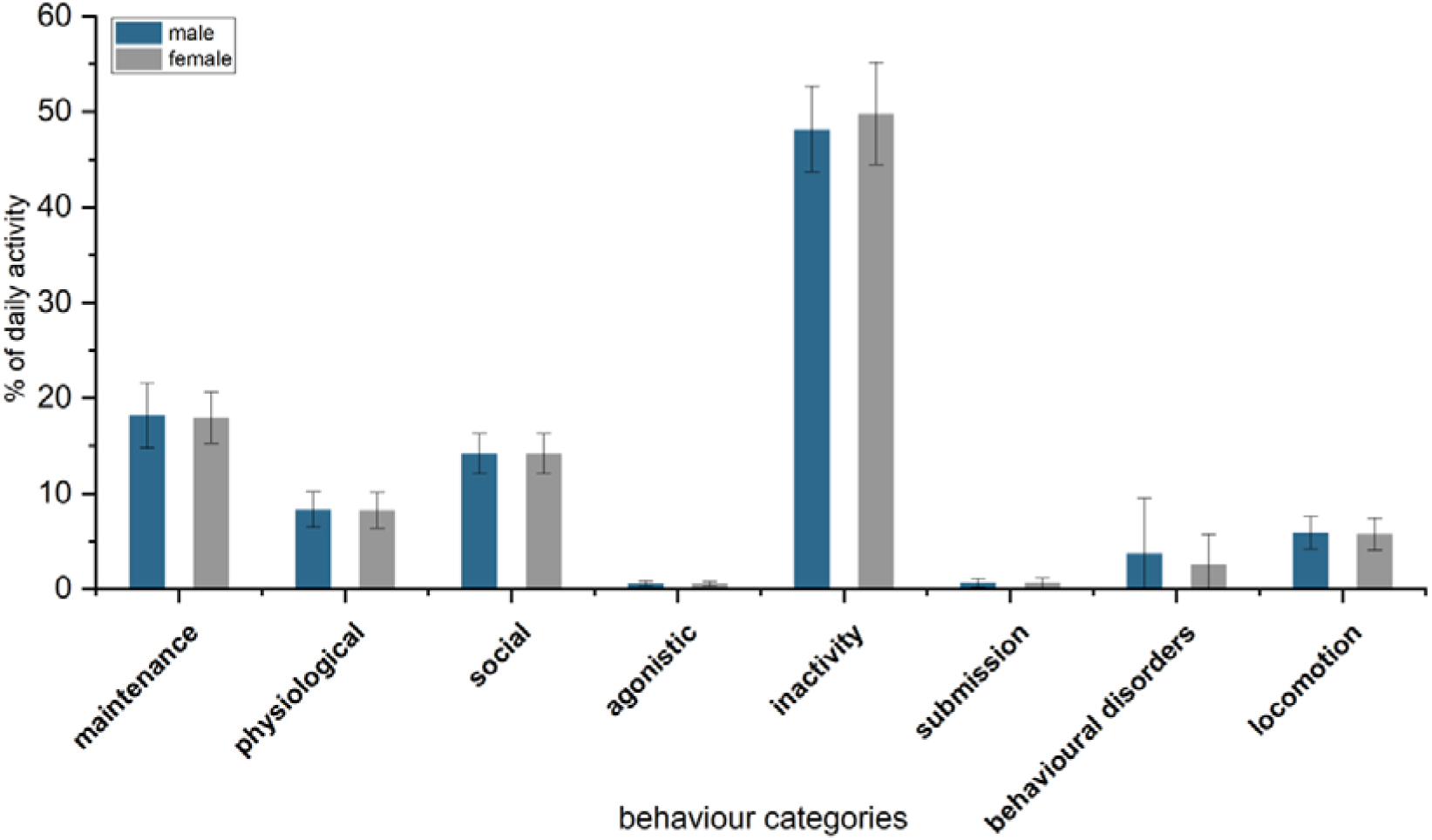
– Time activity budgets of the Spix–s macaw under captive conditions, male (blue) and female (grey).

**Table 1.**
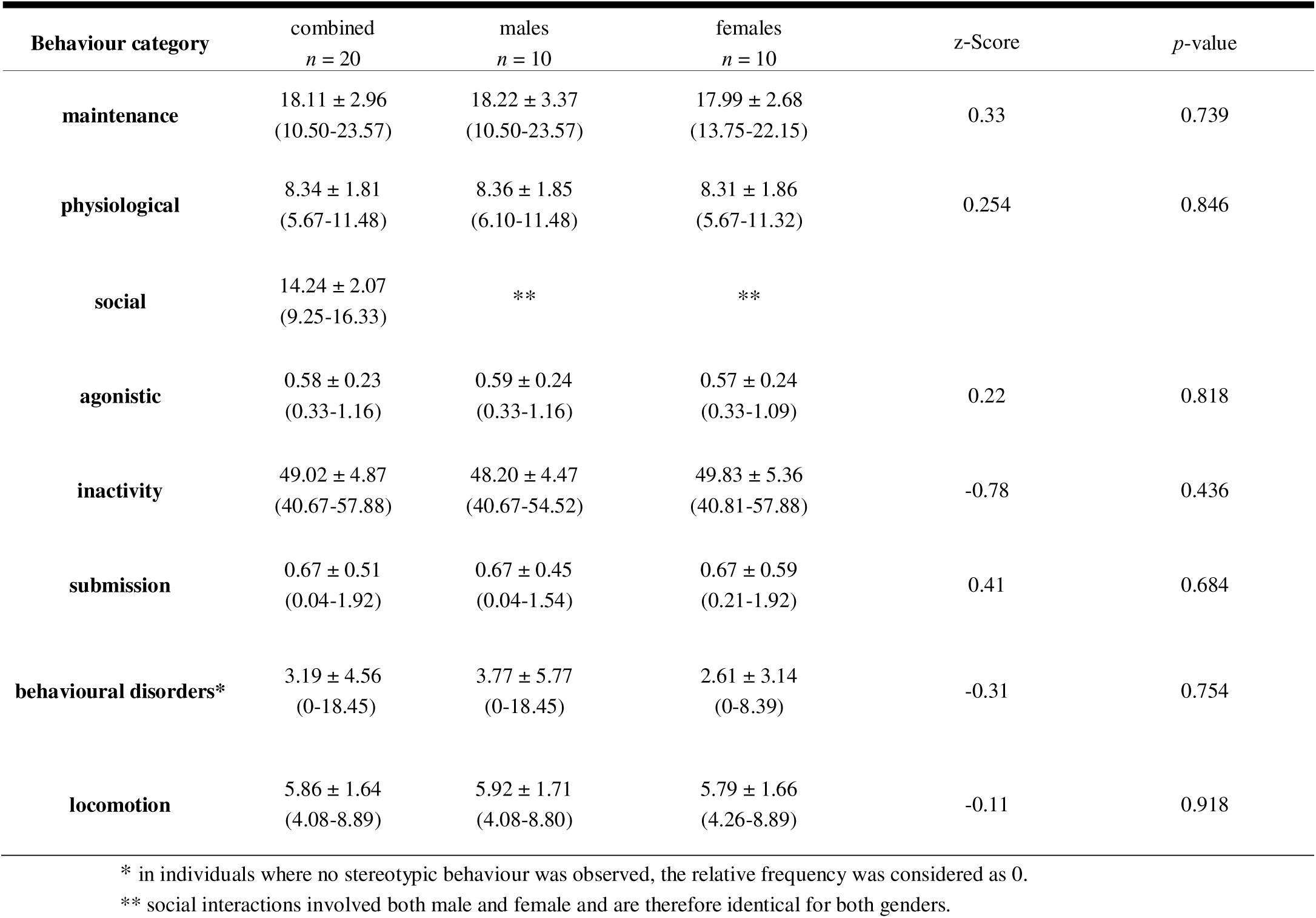
– Summary of activity patterns observed for 20 adult Spix’s macaws and test statistics for the comparison of males and females (exact Mann-Whitney U test). Values for single study groups (combined, male and female) are given as the percentage of the diurnal period spent on each focal activity.

The timing of physiological behaviours followed a bimodal pattern (see Fig. 4), with the highest activity observed between 08:00 and 08:59 and another peak between 16:00 and 16:59. These two peaks were thus closely associated with the feeding schedules. Indeed, foraging accounted for nearly the totality of time spent displaying physiological behaviours. Inactivity and diurnal resting peaked in many individuals during postfeeding periods (10:00–13:00 and 17:00–19:00). Both agonistic and submission behaviour were closely associated with the presence or proximity (i.e., auditory but no visual contact) of the animal keepers. Maintenance behaviours were recorded without any evidence for specific time-activity peaks, although autopreening typically followed prolonged periods of inactivity or allopreening sessions. Locomotion behaviours showed no time-specific pattern, and movement frequency over a given period appeared context-dependent. Behavioural disorders occurred either in association with the direct presence of keepers (as an obvious trigger) or had no identifiable visual or acoustic trigger (often the case in chronic forms of stereotypies). Behavioural disorders were observed in 8 out of 20 observed individuals.

**Figure 4.**
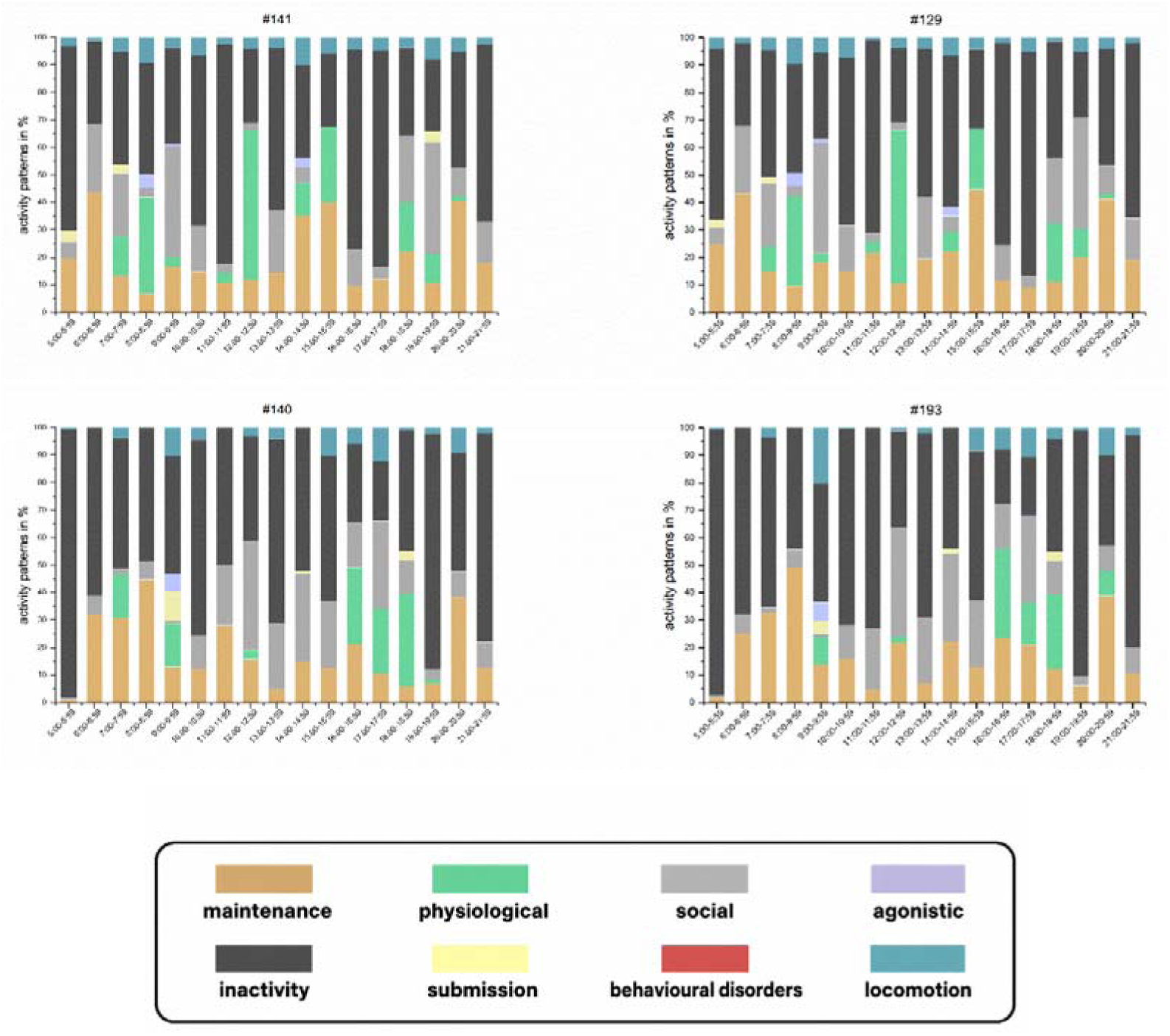
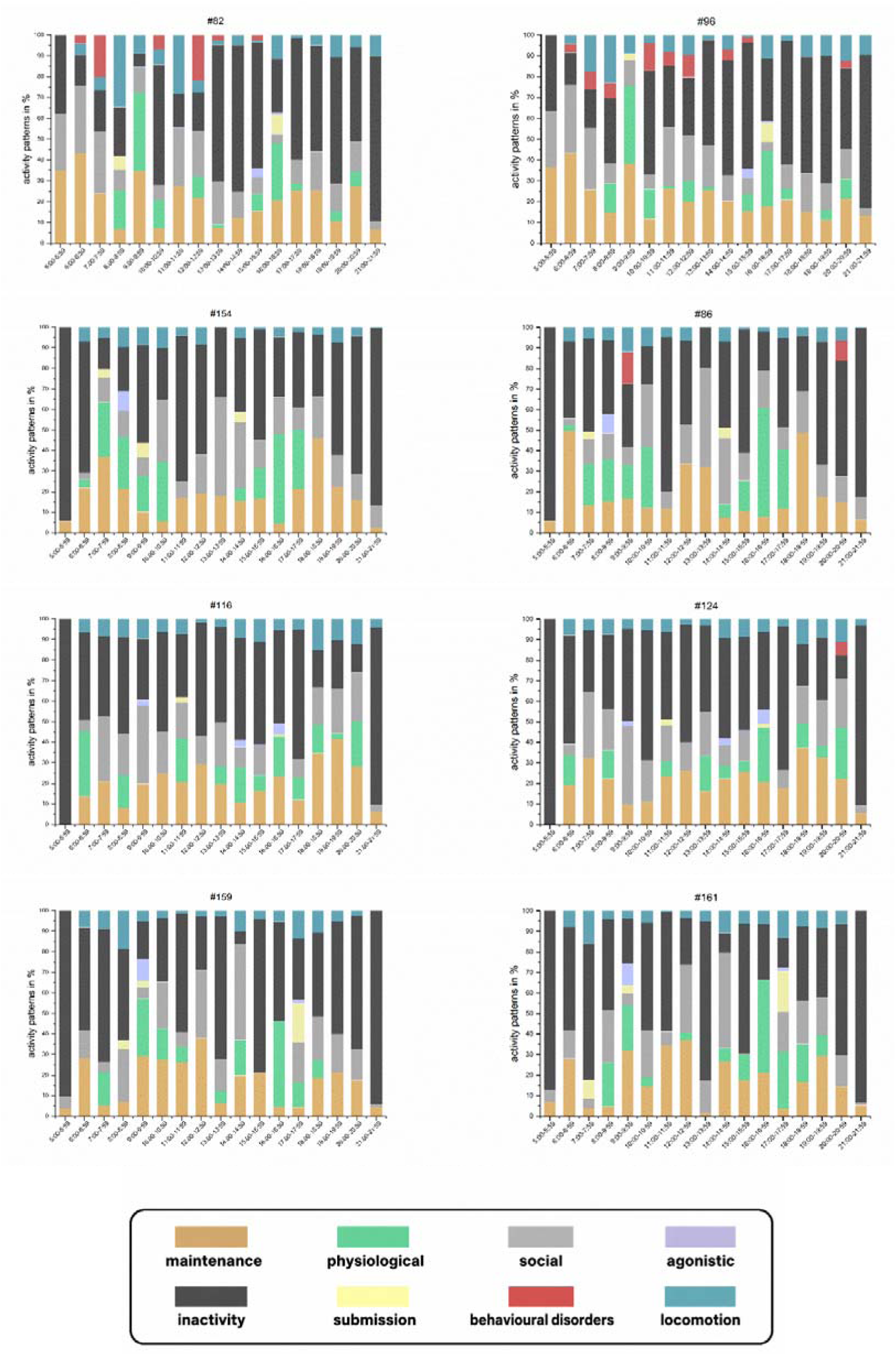
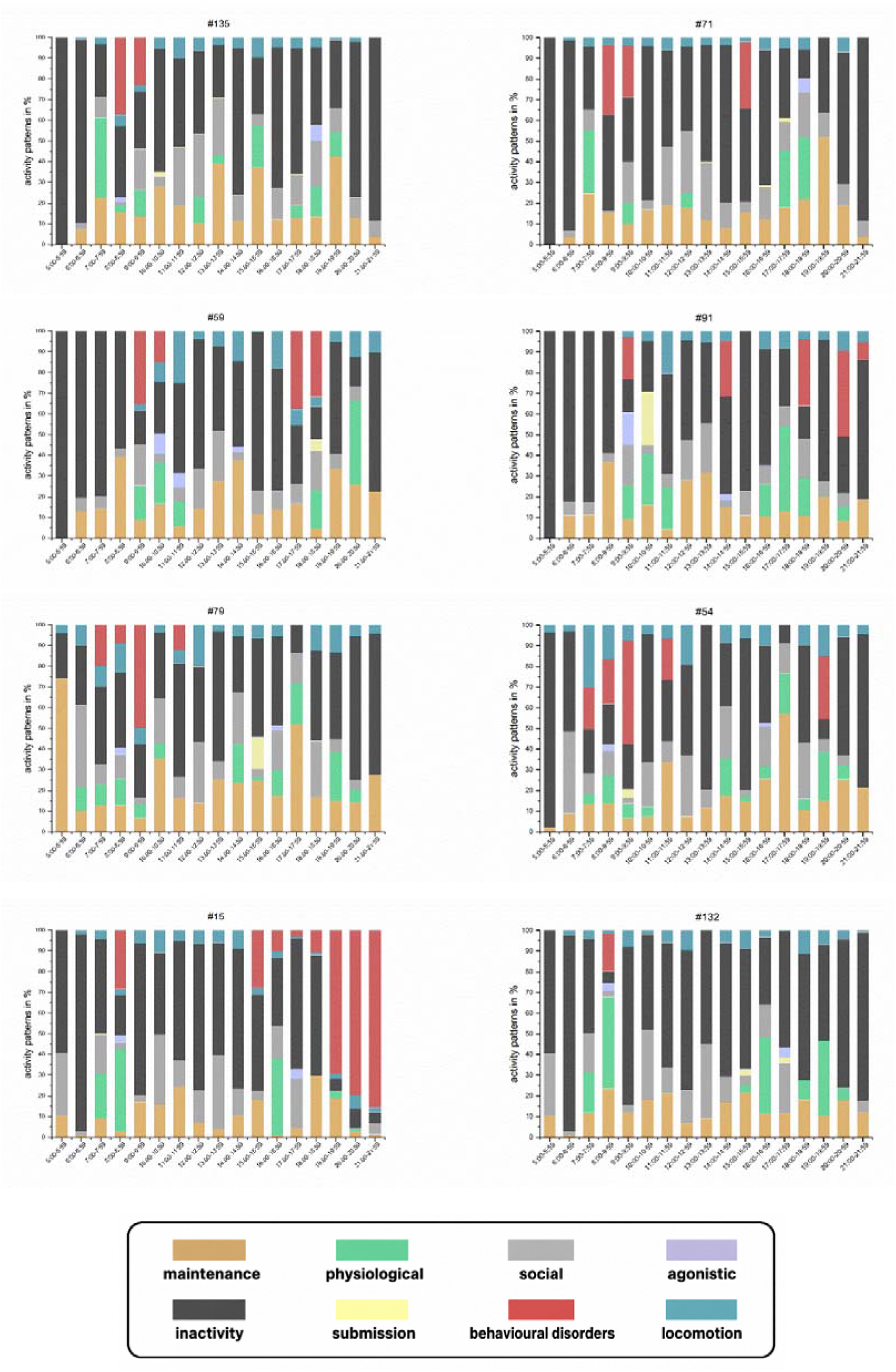
– Diurnal activity patterns of 20 individuals with associated studbook numbers, with each row representing a pair with the male on the left and the female on the right side. Pairs are sorted by decreasing values of intra-pair similarity as measured by the metric *S*. Each row represents one pair. Stacked column bars are presented for males on the left and for females of each pair on the right side, respectively.

### 3.3 Similarity in time-activity pattern and breeding output

The hierarchical cluster analysis demonstrated the presence of intra-pair similarity in time-activity patterns, with five out of 10 pairs being correctly forecasted as actual pairs (see Fig. 5), and three other pairs (#154/#86 – #male studbook ID/#female studbook ID, #141/#129 and #79/#54) showed slight divergences but remained within the same well supported larger clusters (Fig. A10). In contrast, two pairs (#15/#132 and #59/#91) showed a higher divergence in their time-activity patterns. The ANOSIM results confirmed that the 10 defined clusters significantly differ from each other (R = 0.997, p = 0.001).

**Figure 5.**
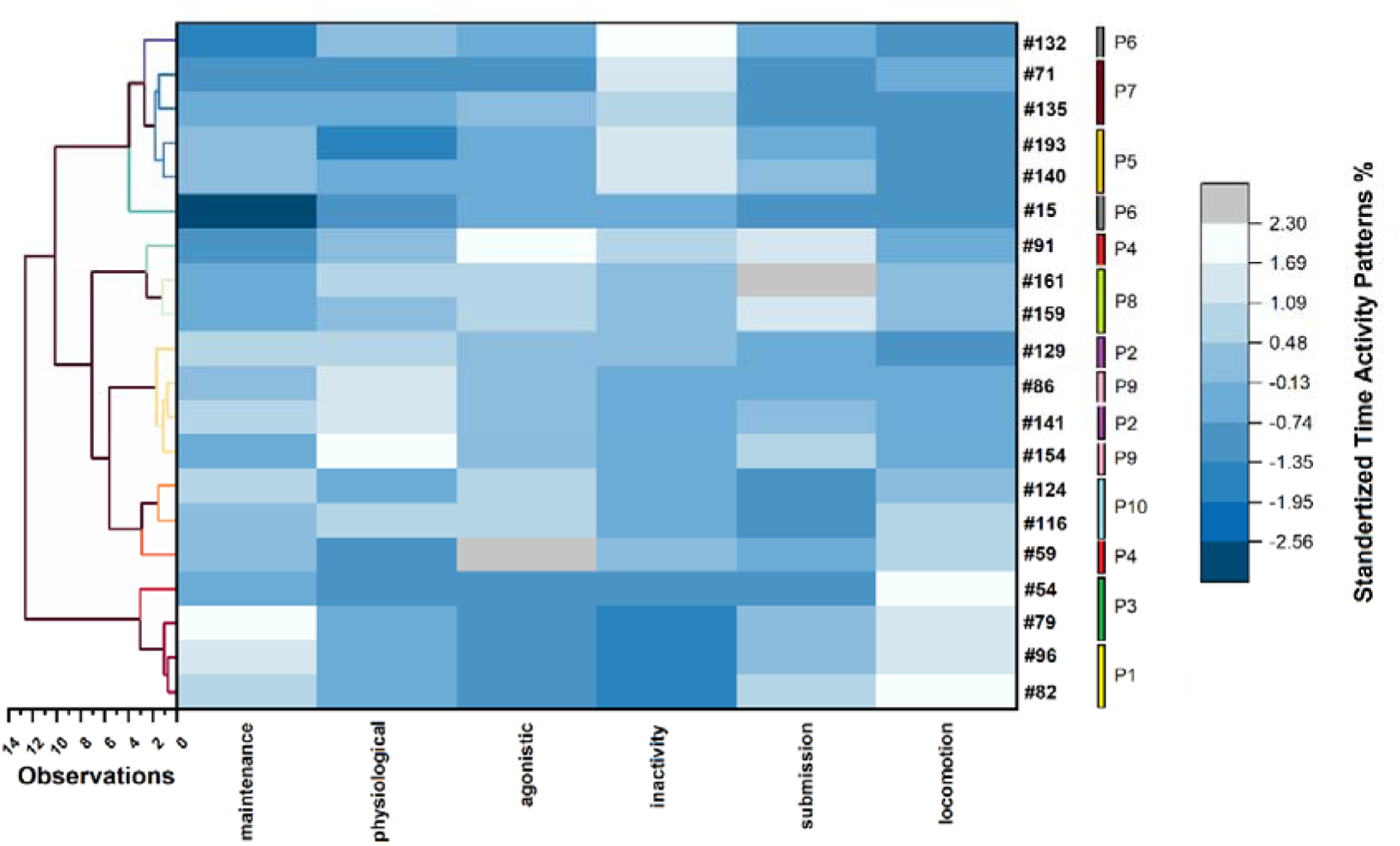
– Heatmap with a hierarchical cluster analysis using standardized data (type = WardD, average type = Euclidean, k = 10 clusters); actual pairs are highlighted (P1-P10) with the associated Studbook numbers, and clusters show the associated individuals with the highest inter-individual similarity. Pairings (with respective male/female) P1: #82/#96, P2: #141/#129, P3: #79/#54, P4: #59/#91, P5: #140/#193, P6: #15/#132, P7: #135/#71, P8: #159/#161, P9: #154/#86, P10: #116/#124.

The similarity index for each pairing resulted in a comparable pattern, with most pairings achieving a *S* value of > 0.75, except for #79/#54 and #15/#132 (Fig. 6). The distribution of time-activity patterns indicates a high intra-pair similarity with a mean □ of 0.78 ± 0.067 across all actual pairs, with an evident overlap between partners in the frequencies and the temporal distribution of the behaviours exhibited (see Fig. 4).

**Figure 6.**
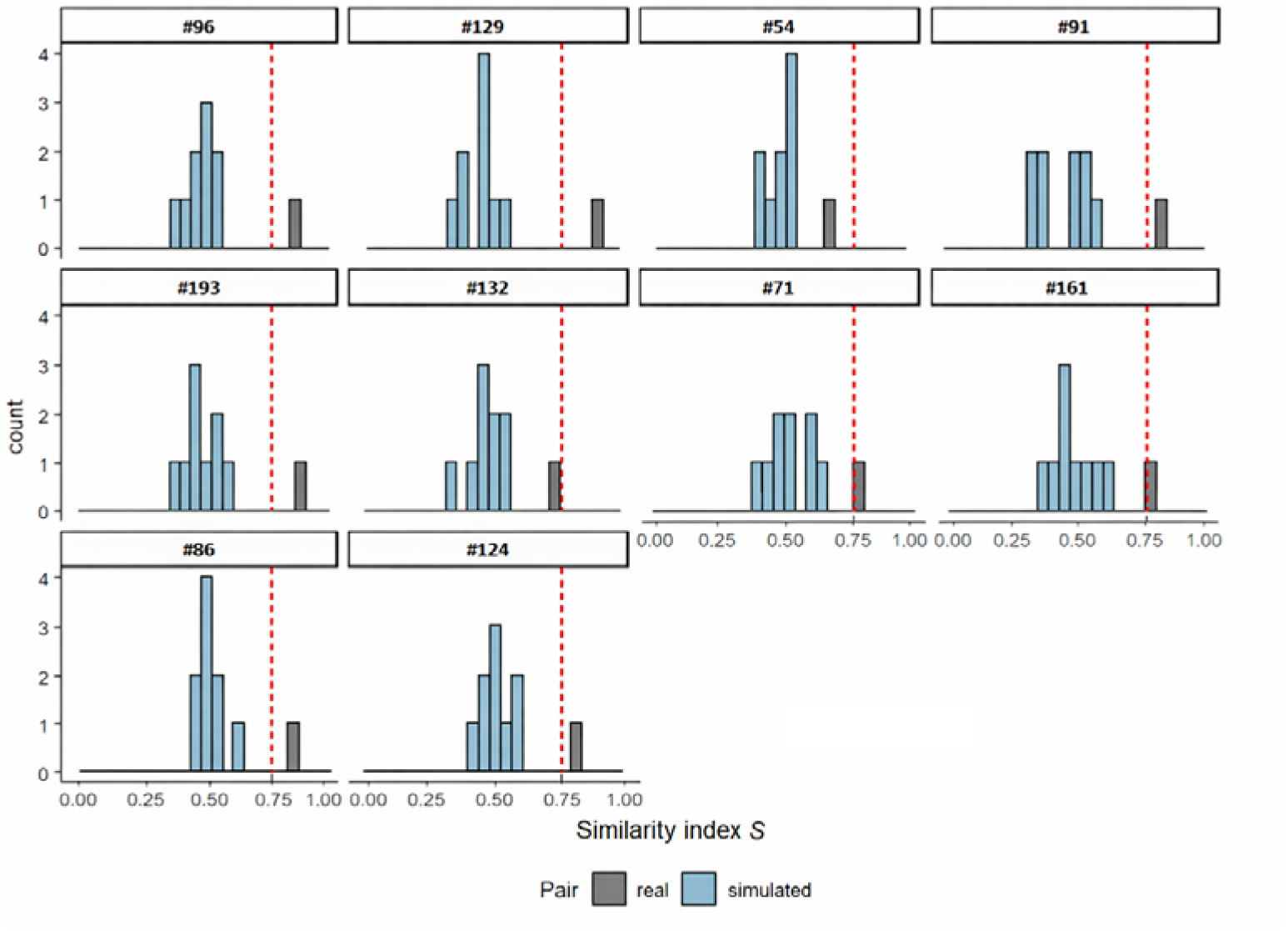
– Similarity indices presented for each female, showing the calculated similarity index for the actual “real” partner in grey (▪) and the simulated partners in blue (▪). The studbook numbers of the females are indicated above each plot. A v-line of *S* = 0.75 is indicated by the red dashed line (---) corresponding to the pairs that were observed to breed.

The simulation of the pairings between each female and all possible males revealed that similarity was always higher with the actual partner than with any other male (exact binomial test, p < 0.0001; Fig. 6).

When linking the similarity index to reproductive performance, we observed that all pairs that bred successfully showed a similarity index higher than 0.75 (Fig. 7). Poor social and behavioural similarity seemed associated with low reproductive performance (Fig. 7), as the pairings #79/#54 and #15/#132 did not produce eggs and were consequently separated in 2019 or 2020, respectively. The pair #59/#91 did not produce offspring at ACTP, but it sired two offspring in 2017 in Qatar, consistent with a similarity index greater than 0.75. An interesting case was the pair #141/#129, which showed the highest similarity index and produced several clutches, yet none of the eggs were fertile. Both #141 and #129 produced offspring with different partners in 2021 (#141) and 2023 (#129).

**Figure 7.**
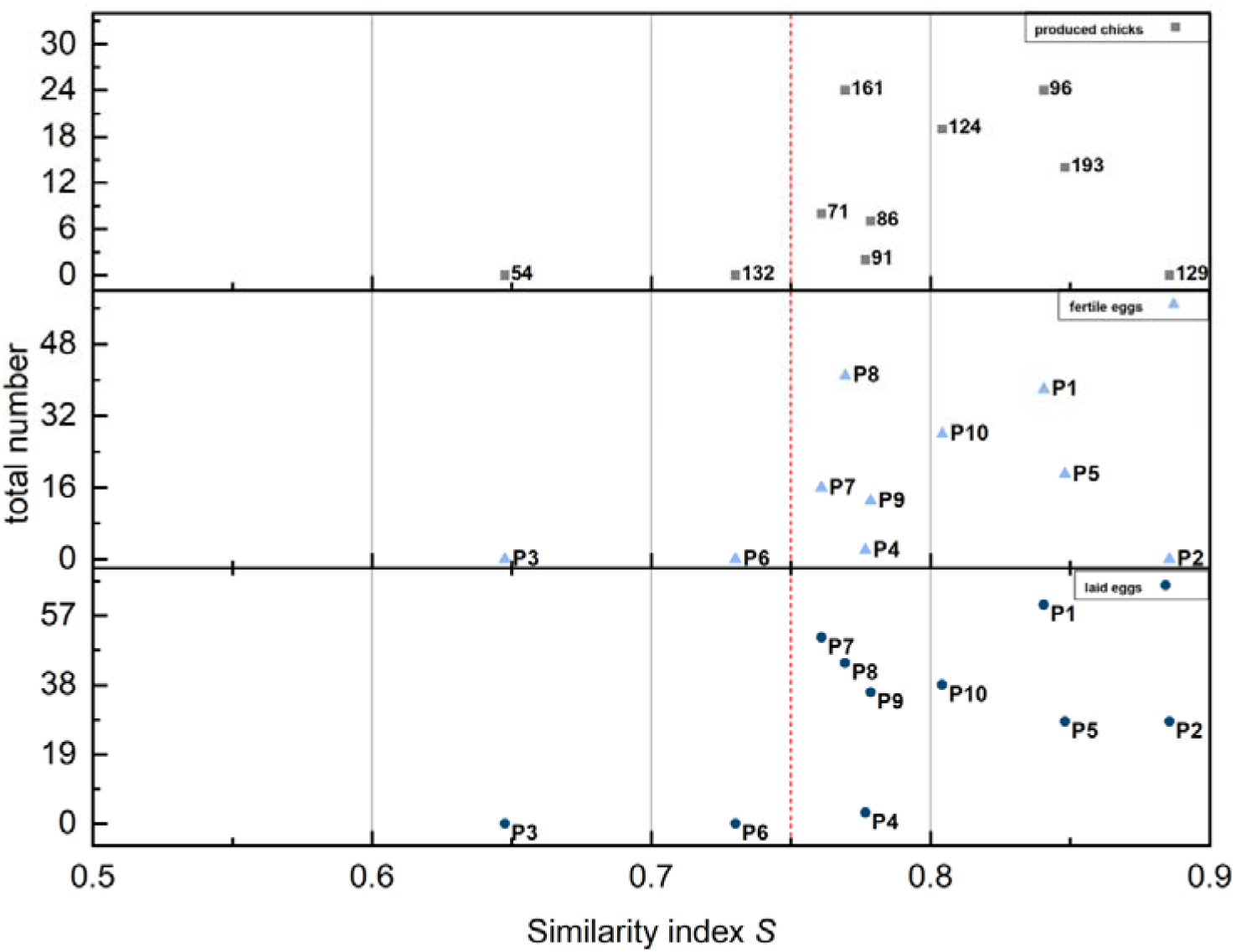
– Scatter plot to illustrate the relationship between the Similarity index (*S*) of the observed ten pairings and the total number of laid eggs (⍰), fertile eggs (▲), and chicks (▪) produced between 2018–2024. The red dashed line (---) indicates the similarity index *S* = 0.75 corresponding to pairs observed to have bred. The IDs of individuals forming the respective pairs are provided in the legend of Fig. 5.

Our experience is that established pairs often showed increased behavioural synchronization within the first six weeks of pairing, which allowed ACTP staff to assess pairing success in time before egg laying. Using this information to improve the compatibility between pairs, together with nutritional improvement and environmental enrichment, resulted in a steady increase in offspring numbers at ACTP Germany: 2019 yielded 11 offspring, 2020 yielded 21, and 2021 yielded 51. Up to four clutches were laid by each pair; we restricted this to two for the years from 2022 onwards. This yielded 35 offspring in 2022, 42 offspring in 2023, and 44 offspring in 2024. It also increased the number of parents capable of successfully raising their chicks from 0 in 2019 & 2020 to 2 in 2021, 3 in 2022, 7 in 2023, and 16 in 2024. There was a similar increase in fertility rates, starting with 27% in 2019, 30% in 2020 when several young pairs laid their first clutch, 47% in 2021, 45% in 2022, 49% in 2023, and 64% in 2024. These results are based on a total of 240 produced clutches recorded (26 in 2019, 49 in 2020, 53 in 2021, 37 in 2022, 44 in 2023 and 31 in 2024) and 86 forced mating attempts conducted during the same time period at ACTP Germany (25 in 2019, 14 in 2020, 16 in 2021, 8 in 2022, 14 in 2023 and 9 in 2024).

## 4. Discussion

Our main objectives for this study were to (1) describe the full suite of behaviours of Spix’s macaws in human care, (2) document their time-activity patterns, and (3) investigate the degree of intra-pair similarity in time-activity patterns and its relation to breeding performance. We now discuss the results in view of potential applications in the management of *ex-situ* breeding programs as well as for the reintroduction of parrots into the wild.

### Ethological data, behavioural disorders and implications for animal welfare and conservation

Our study provides the first ethogram for the Spix’s macaw, comprising a total of 85 behaviours, including stereotypies and other behavioural disorders (hereafter, *behavioural disorders*). These complement ethograms previously published for various Old World and New World parrots, including *Eupsittula canicularis* (Hardy, 1963), *Agapornis* spp. (Dilger, 1960), *Calyptorhynchus lathami* (Pepper, 1996), *Trichoglossus* spp. (Serpell, 1979), *Loriculus* spp. (Buckley, 1968), *Cyanoramphus* spp. (Higgins, 1999), *Nestor notabilis* (Keller, 1976), *Cacatua* spp. (Noske *et al.,* 1982; Higgins, 1999; Rowley, 1990), *Amazona* spp. (Levinson, 1980; Lantermann 1987; Snyder *et al.,* 1987; Prestes, 1991; Queiroz *et al.,* 2014) and other neotropical species (Ayres-Peres and da Silva, 2017).

Most of the behaviours of Spix’s macaw appear to be similar to those recorded for closely related species in both captive and wild environments (Uribe, 1982; Christiansen and Pitter, 1992; Pitter and Christiansen, 1995; 1997; Schneider *et al.,* 2006; Favoretto *et al.,* 2024). We recorded a total of 12 behavioural disorders. The stereotypies, which we classified as physical (6-10 in Appendix I 3.1.9), have also been reported for other parrots (Luescher, 2006; Acharya and Rault, 2017). For example, feather plucking—a known problem in Spix’s macaw populations (Hammer and Watson, 2012)—is ubiquitous in captive stocks (van Zeeland *et al*., 2009). In contrast, other stereotypical displays we observed are little discussed in the literature (e.g., non-physical stereotypic displays, infanticide), which could imply that some of these behaviours are species-specific, or that they are little studied, or both. While no wild parrot has been documented to exhibit the aforementioned disorders, behaviours we consider here to be disorders in captivity may occur in nature with a different aetiology, such as egg destruction and infanticide (see Heinsohn *et al*., 2011). Such behaviours may occur in the wild in the context of intraspecific or interspecific competition. In our study population, it was observed only in some individuals, whereas others were never observed to engage in such behaviour.

In terms of time-activity patterns, resting was the predominant behaviour, followed by maintenance and social behaviour as observed in other captive parrots such as Scarlet macaws (*Ara macao*, Cornejo *et al.,* 2005), Lear’s macaws (*Anodorhynchus leari,* de Azevedo *et al*., 2016), Hyacinth macaws (*Anodorhynchus hyacinthinus*, Checon *et al*., 2020), Vinaceous-breasted amazons (*Amazona vinacea*, Ramos *et al*., 2020) or Senegal parrots (*Poicephalus senegalus*; Lantermann, 1998). While there is no such data for wild Spix’s macaws, studies on other species suggest that a prolonged period of inactivity is a hallmark of captivity. For example, around a year after their release, Scarlet macaws spent 35% of their time resting compared with 41% for conspecifics in human care (Cornejo *et al.,* 2005).

We recorded Spix’s macaws to spend 8.3 ± 1.8% (5.7-12.4) of their full diurnal activity period foraging, which is lower than the value of 15% reported for captive Scarlet macaws (Cornejo *et al.,* 2005), but comparable to estimates provided for Hyacinth, Scarlet and Military macaws (*Ara militaris*) from the Loro Parque Zoo (Britsch, 2018). Although foraging activities vary markedly between individuals and environments, foraging activities were reduced in captivity. In their natural environment, parrots spent a substantial amount of time foraging, as demonstrated for released Scarlet macaws (28%; Cornejo *et al.,* 2005), wild Ouvéa parakeets (*Eunymphicus uvaeensis*) (47%; Robinet *et al.,* 2003), or wild Glossy black cockatoos (*Calyptorhynchus lathami halmaturinus*) on Kangaroo Island (26% for non-breeding, 36% for breeding birds; Chapman and Paton, 2005). Such divergence between wild and captive birds is expected, given that foraging activities include search flights to locate feeding sites, handling food items, and interactions with competitors for access to resources (Chapman and Paton, 2005; Brightsmith *et al*., 2018). While comparative data on locomotion are scarce, we observed Spix’s macaws spending even less time actively moving than during foraging, which is likely a response to easy access to food and the captive environment.

By eliminating some constraints, such as unpredictable food distribution and availability, competition, and predation, conditions in human care induce a shift in activity patterns that may promote the expression of behavioural disorders. Compared to its (presumed) absence in the wild, we observed Spix’s macaws to spend, on average, 3.2% of their full diurnal activity period exhibiting behavioural disorders. The time budget dedicated to such behaviours varied widely among individuals—ranging from completely absent in some to up to 3 hours per day for one individual (#15). The exact aetiology of behavioural disorders remains unclear. In ACTP’s captive population, it probably results from the lack of some activities, possibly the lack of social interactions and environmental challenges, in conjunction with stress factors resulting from husbandry conditions initially suboptimal. The latter comprises inappropriate hand-rearing, lack of enrichment, and poor health management, but other intrinsic factors may also play a role, such as personality, stress levels, or genetics (see Garner *et al*., 2006; Luescher, 2006; Owen and Lane, 2006; Cussen and Mench, 2015).

Reducing the occurrence of behavioural disorders by optimising husbandry protocols is an important way to promote animal welfare and the productivity of *ex-situ* populations. This is because such behaviours can lead to physical injuries by auto-mutilation, redirected aggression, and/or feather plucking (Owen and Lane, 2006; Luescher, 2006; Acharya and Rault, 2020). Behavioural disorders in captive parrots can also interfere with breeding. As an extreme illustration, ACTP staff observed a few females purposefully destroying their eggs during the incubation period or killing their offspring after hatching.

Husbandry should therefore aim at creating an environment that minimises the occurrence of behavioural disorders (Coulton *et al*., 1997; Field and Thomas, 2000; Meehan *et al.,* 2004; Wang *et al*., 2009; van Zeeland *et al*., 2013; Reimer *et al*., 2016; Rodriguez-Lopez, 2016; de Almeida *et al.,* 2018; Livingstone, 2018). For this purpose, behavioural data serve as an important template to improve husbandry guidelines. Several measures were taken to reduce behavioural disorders in ACTP facilities following data collection and analysis. For example, the environment of all birds has been frequently enriched since the behavioural data collection for this study was completed, by providing them with paper rolls, cardboard, new modular toys, treating dispenser toys, and fresh greens. ACTP staff have also progressively favoured the rearing of chicks by their parents. We have yet to investigate and disentangle the effects of such changes on time-activity patterns, but behavioural disorders in ACTP facilities have substantially dropped over time. We assume that minimising the occurrence of behavioural disorders will improve the success of conservation breeding and the success of reintroducing animals to the wild. Indeed, the expression of behavioural disorders can impede an individual’s ability to respond adequately to environmental changes and limit the capacity to learn or develop behavioural strategies that are important for survival. In addition to genetic criteria (inbreeding, relatedness) and the bird’s physical condition, we retained only individuals showing no evidence of behavioural disorders for reintroduction. The first two cohorts selected in this way were therefore released into the wild in June and December 2022, and the first wildborn offspring successfully fledged in May 2024 (Purchase *et al*., 2024; Vercillo *et al*., 2024). While these results are encouraging, more effort could be directed toward bringing the behavioural profiles of captive birds closer to those of wild ones. For example, the time-activity pattern of wild psittacines is influenced by temporal changes (e.g., Chapman and Paton, 2005), so modifying the captive environment to induce similar changes may prove beneficial.

### Behavioural similarity and implications for conservation breeding efficiency

Another way to increase the productivity of animals in captivity is to provide individuals with partners with whom they are willing to mate (Martin-Wintle *et al.,* 2015; *et al.,* 2019; Alverson *et al*., 2023). This can be achieved in three main ways. One solution is to let animals freely choose their partners as they would do in the wild. Free mate choice has indeed been linked to higher breeding output in psittacines (Waugh and Romero, 2000; Luescher, 2006; Spoon *et al*., 2007), mammals (Parrott *et al*., 2019) and reptiles (Lemm and Martin, 2023). Letting animals choose their mates freely is usually not feasible in captivity due to time and space constraints or limited availability of potential partners. Moreover, free mate choice may not always result in the desired conservation outcome of maintaining high genetic diversity. A second possibility is to expose, in a controlled setting where actual copulation is impossible, a focal individual to a few candidate mates and infer its mate preferences by recording its behavioural responses. While this has proved successful in some species (Martin-Wintle *et al.,* 2015; Alverson *et al.,* 2023), assessing mate preferences in this way requires a large, highly specific enclosure layout. Finally, breeders may also adjust pairings based on inferred behavioural compatibility (Spoon *et al*., 2007; Fox and Millam, 2014). Breeders tend to consider members of pairs which spend a lot of time together as “harmonious pairs” and a harbinger of good productivity. Indirect evidence suggests that breeders’ empirical knowledge may be correct. For example, in cockatiels (*Nymphicus hollandicus*), more eggs were laid, and more chicks hatched and were reared by mates that were more synchronised, spent more time in close distance to each other, exhibited frequent allopreening, and displayed fewer aggressive acts (Spoon *et al*., 2007). Raw behavioural data could therefore be helpful for evaluating intrapair “harmony” and forecasting the chance of successful breeding. Unfortunately, how breeders assess such behavioural compatibility in practice when mate choice is constrained has been little explored. Our approach was to design a similarity metric (*S*) to compare the time-activity patterns of each pair of individuals and to investigate whether this metric predicted breeding performance.

Our results are consistent with the view that behavioural compatibility may be an important factor shaping the reproductive output of parrots. In our limited data on Spix’s macaws, the two pairs with the lowest behavioural similarity did not breed, whereas all but one pair with high similarity values did. The single pair with a high similarity value that did not produce any offspring (#141/#129) did produce many eggs. A pathological cause seems unlikely to explain the reproductive failure of this pair, as both #141 and #129 later produced offspring with different partners. A poor genetic match that led to the expression of lethal alleles might be a possible explanation, given that the entire population is highly inbred and based on only six founder individuals (Purchase, 2019). Another pair (#59/#91) also showed poor breeding performance and a reasonably high level of behavioural similarity. It did not produce offspring at ACTP Germany, although it sired two offspring in 2017 in a facility in Al Wabra Wildlife Preservation, Qatar (#216 and #217, Purchase, 2019).

Our sample size was arguably small, and further data should be collected on Spix’s macaws and other species to assess the usefulness of the similarity index and alternative metrics used to measure behavioural compatibility in conservation breeding practice. For the time being, our results and practical experience suggest that the breeding output of forced pairing may be improved by selecting candidate mates using behavioural data, in addition to other criteria that may be employed, including age, genetics, and previous breeding success. Assuming that behavioural compatibility is a good predictor of breeding output, another question remains: when would measuring behavioural compatibility provide the best forecast of that breeding output? It seems tempting and practical to measure behavioural similarity on isolated individuals. However, time-activity patterns are influenced by the social environment. We indeed observed that all females showed the highest similarity with their actual partner in comparison with other males, which is not surprising given that the Spix’s macaw is a gregarious species that tends to form temporary bonds and flocks with conspecifics, aligning its behaviour within the flock (see Hobson *et al*., 2014). For these reasons, we suggest that behavioural similarity should be evaluated after placing a female and a male together for a fixed period of time. Our experience is that established pairs often show increased behavioural synchronisation within the first six weeks of pairing, so this duration seems appropriate. Measurements should be taken before the onset of the breeding season to allow sufficient time to adjust pairs before egg laying. Ideally, this should not be done too long before the breeding season in case mate preferences vary over time.

Whenever they detect poor similarity, ACTP staff now either swap pairs immediately when similarity was really low or do so at the onset of the next breeding season when similarity is intermediate. Although not always performed quantitatively as in this paper (due to time constraints), it was also sometimes based on the subjective perception of the staff; re-pairing based on similarity helped reevaluate and adjust >90% of all pairings between 2019 and 2024 that had either never bred or usually produced infertile clutches. The facility’s reproductive output has increased markedly since the practice was implemented. This practice, however, was initiated alongside other general improvements in husbandry conditions (nutritional improvements and environmental enrichment), making it difficult to infer causality. To increase breeding performance even further, groups of six juveniles (3 males and 3 females with low relatedness) have been placed together in enriched and enlarged aviaries since 2023. This allowed them to live together until they freely choose their partner, which occurred around 3 yrs old. After this age, which coincides with the appearance of territorial behaviour, the current plan is to place each pair in a separate breeding aviary. The future will show whether this is the appropriate way forward.

## 5. Conclusions

Our study highlights that detailed monitoring of behaviour can benefit ongoing or planned *ex-situ* programs, especially for socially complex species such as the Spix’s macaw and many other parrots. Behavioural studies can help identify welfare issues, including stereotypies, other behavioural disturbances, and displacement behaviour, and improve husbandry guidelines to enhance the effectiveness of *ex-situ* breeding programs. The discovery of behavioural disorders and prolonged inactivity among Spix’s macaws in human care at ACTP Germany prompted ACTP Germany to enrich the environment and modify breeding protocols. The finding that behavioural similarity between paired individuals could reliably predict breeding performance also prompted them to revise the procedures for matching partners based on behavioural criteria. We documented here how breeding productivity improved by combining these changes with other improvements in husbandry conditions. Importantly, the relevance of behavioural monitoring extends beyond the management of *ex-situ* breeding programs. Characterising individual behaviour is also key to selecting candidates for release into the wild and improving the success of reintroduction programs. In particular, ethograms are necessary to assess whether selected birds successfully adjust to their novel environment (see Purchase *et al*., 2024; Vercillo *et al*., 2024).

While behaviour is not the only factor determining animal welfare and breeding productivity, considering this information can improve conservation breeding in practice. We would like to end by urging others to participate in the global effort to integrate behavioural studies into conservation science (Curio, 1996; Buchholz, 2007; Berger-Tal and Saltz, 2016; Snijders *et al*., 2017). The strive to improve breeding conditions within any single breeding facility makes it difficult to disentangle the effects of many concurrent decisions. Only the combination of findings from facilities with distinct husbandry setups and focal species will yield a robust scientific body of knowledge to guide *ex-situ* conservation.

## Author contributions

Conceptualization, V.M., A.C., H.H.; Data curation, V.M., A.C., Methodology, V.M., A.C., K.S., J.A.; Software, V.M., A.C.; Formal analysis, V.M., A.C.; Investigation, V.M., K.S., J.A.; Data curation, V.M.; Writing – original draft preparation, V.M.; Writing – review & editing, V.M., A.C., P.S.C., C.P., L.C., H.H.; Visualization, V.M.; Supervision, L.C., A.C., H.H.

## Data Availability Statement

The data presented in this study are available upon request from the corresponding author. The data is not publicly available due to the internal policy of ACTP.

## Acknowledgments

We would like to thank Mr. Martin Guth for allowing us to conduct the research and for providing funding for the equipment. We would also like to thank the Akademie für Vogelkunde and the Vereinigung für Artenschutz, and the Vogelhaltung und Vogelzucht e.V. for funding the PhD studies of VM. We thank all the institutions that supported ACTP, including ProCom/Grumbach, NuTropica®, Birdfarm.de, Kauri CAB, AVES publishing, Bowman Books, Al Wabra Wildlife Preservation, Parrots International and Naturkunde Museum in Berlin. We would like to thank Michał M. Hryciuk, Brij K. Gupta and two anonymous reviewers for their comments on the early draft of the manuscript.

