## Appendix I for "The value of behavioural activity records for conservation breeding: the case of Spix’s macaw in human care"

**Contents**

(1) Detailed methods
(2) Appendix I: Ethogram
    a. Maintenance behaviours
    b. Physiological behaviours
    c. Locomotion behaviours
    d. Inactivity behaviours
    e. Agonistic behaviours
    f. Displacement behaviours
    g. Submission behaviours
    h. Social behaviour
    i. Sexual behaviour
    j. Behavioural disorders
(3) Appendix I: Comments
(4) Appendix I: Additional references
(5) Appendix I: Illustrations of behaviours
(6) Appendix II: Bootstrap analysis

The following supplementary material presents all behaviours identified and their classification into different categories, which we used to establish the ethogram and to compile time-activity patterns.

### (1) Detailed methods

*Observations.* To define and describe individual behaviours and compile the ethogram, birds were observed weekly in 2018 and 2019, either directly during the facility's operational activities or through the indoor cameras.

Documentation of the entire suite of behaviours (qualitative ethograms) was based on observations of 123 individuals, including 29 breeding pairs as well as six groups of individuals scheduled to be part of the first release in the wild. These six groups included four small groups (5, 6, 7 and 8 individuals, respectively) and two large ones (18 and 21 individuals, respectively). Non-contact observations were carried out using cameras to avoid behavioural changes influenced by the presence of an observer near the aviary. In addition, the behaviour of released individuals was observed during three weeks in July 2022 (*n* = 8) and August 2023 (*n* = 13) using binoculars (10 x 40) during the peak activity time period from 05:00–07:30 and 15:30–18:00. We describe a total of 50 novel behaviours not covered by Marcuk et al. 2020.

*Data analysis*. Behavioural durations were determined using video frame analysis in Avidemux (v. 2.7.4). All descriptive statistics are given as mean ± standard deviation (SD), with the range in parentheses. To compare the predominance of leg use (laterality) between sexes, we used a proportion test. To determine whether the right or left leg is preferred, we used a binomial test. All statistical tests were performed in R (v. 4.3.4, R Core Team, 2024) with a significance level of *a* = 0.05.

### (2) Ethogram

#### a. Maintenance behaviours

This behaviour category is defined as being directly or peripherally associated with body and plumage maintenance. The individual behaviour patterns are described as follows.

1. **Body shake** (Figure A1a): An accelerated shaking movement with an entirely fluffed plumage, with wings moving away from the body, while remaining partly unfolded. Usually followed by an allopreening or auto-preening session or occurring after a prolonged period of inactivity. In comparison to the displacement form (see below), body shaking is not accompanied by submissive calls or eye-blazing. The behaviour seems intended to rearrange the wing and upper-body plumage feathers or to remove dust from them. Body shakes lasted on average 0.60 ± 0.20 s (0.4–1.1; *n* = 16), often consisting of multiple singular shakes that last about 0.2 s.

2. **Head** **scratch** (Figure A1b): Rapid, alternating movement of the front claws, while the head remains in a slightly lowered position and is directed towards the focal leg. Head scratching seems intended to function as a controlled removal of any type of foreign matter (scales, dust, food remains) from the head and neck region. This behaviour is most frequent during the post-feeding period or following allopreening, involving very rapid, consecutive movements of the toes. Scratching varied in duration; the analyzed sequences lasted, on average, 3.35 ± 1.35 s (0.76–8.17; *n* = 24).

3. **Head shake** (Figure S1c): Repetitive head flicks from one side to another, performed perpendicular to the body alignment. Single head shakes lasted between 0.1 and 0.2 seconds. Mean duration was 0.29 ± 0.09 s (0.17–0.47; *n* = 20).

4. **Tail wag**: Rapid, bilateral movement of the tail to invoke a rearrangement of the rectrices. Observed often as an element during preening duties or performed during the post-bath period.

5. **Wing & leg stretch** (Figure A2a): Simultaneous, sideward and backward directed stretching of one wing and the collateral leg away from the body, with a time-delayed fanning of the tail feathers towards the same direction. Either followed by bilateral wing stretch (see below) or performed solely or repeatedly for both sides. The average duration for this behavioural sequence was 8.44 ± 1.86 s (5.07–13.71; *n* = 100). Observed most frequently during dawn or after a prolonged period of inactivity.

6. **Bilateral wing stretch** (Figure A2b): Simultaneous stretching of both wings either partly, with carpal joints raised over the back without touching each other, or wings fully extended toward the dorsal direction. Thereafter, relaxation of both wings in their resting position. The full behaviour sequence lasted on average 1.99 ± 0.39 s (1.30–2.87; *n* = 88). The functional context is the same as that of the wing and leg stretch (co-functional behaviours).

7. **Yawn** (Figure A2c): Extension of upper and lower beak under maximal contraction of the mandibular muscles, with head slightly withdrawn backwards. This behaviour was most frequently observed after prolonged inactivity (87.76% of observations; *n* = 144). This behaviour can be occasionally induced if the area between the auricular patch and anterior throat part is mechanically stimulated.

8. **Bill grind**: Repeated bilateral, mutual rubbing of the upper and lower beak. An auto-maintenance behaviour that is often associated with the post-feeding period. This behaviour can also take the form of a displacement behaviour element, occurring in response to the direct presence of an intruder and directed towards the opponent.

9. **Bill wipe**: Rubbing of the beak on a solid surface (perch) first on one side, then repeated for the other; habitually repeated multiple times for both sides to remove any remaining food particles.

10. **Touch-foot** (Figure A3a): Forms a part of the auto-maintenance behaviour, where the head is lowered vertically, and the focal leg is moved towards the head.

11. **Auto-preening** (Figure A3b-d): Auto-preening describes a multi-functional behaviour complex composed of several distinct behaviours (tail preening, Figure A3c, breast preening, back preening, Figure A3b, and wing preening, Figure 3d). The purpose of these behaviours is likely to ensure the maintenance of different plumage regions and removal of flaking scales or feather dust. Preening implies the coordination of tongue and beak to mechanically extract foreign matter from single feathers or carefully extract feathers from the body by repeatedly grasping the respective feathers and working along the barb towards the calamus with rapid movements of the tongue.

12. **Bath**: Bathing occurred at irregular intervals (usually after a long period of inactivity), even if water of sufficient quality and temperatures were above 20 °C. Spix's macaws prefer to bathe during the local midday period (when the sun is nearly at its zenith) and use either a bowl filled with fresh water or an artificial rain system, with the latter clearly preferred (17 vs. 76 documented records). If bathing in a bowl, the bird immerses its head and beak in the water while remaining perched close to the bowl's edge. The head is dipped a few centimeters below the water surface sharply and twisted either sideways or backwards to splash water over the plumage while remaining underwater. This behaviour is often followed by dragging the breast (breast-drag), with the intention of dragging the upper body in a bilateral direction; from one side to the other under the water surface; and/or forcefully dunking wings (wing-dunk) and the upper body into the water if bowl size allows for this. The latter two elements are achieved through a body shake, which redistributes water across all body regions. Head-scratching, tail-wagging, and auto-preening were among the most frequent behavioural elements observed during the post-bathing period. Recorded bathing sessions lasted 643.78 ± 102.2 min (452–668, *n* = 9).

When the artificial rain system was used, individuals showered by hanging in the vicinity of, or directly below, the water sources (nozzles; Figure A4). This behaviour is performed either continuously or in an interrupted manner, then resumed. Birds use different approaches to shower while hanging from the mesh: (i.) Beak-hang (Figure S4c): Performing individual hangs vertically, grasping the mesh only with the beak. The wings and legs are partly unfolded and hang down. (ii.) Backside-wing flip (Figure A4a): The individual hangs upside down using both legs; the wings remain fully extended and are flapped repeatedly; the head is slightly moved backwards, and the back is pointed out towards the water nozzles. (iii.) Upside-hang (Figure A4b): The bird uses either one or both legs, which are passed under the wings and positioned next to the head. The body is either aligned vertically or slightly diagonally (especially when only one leg is involved). Bird hangs using the front claws; (iv.) Umbrella posture (Figure A4d): like backside-wing flip but performing individuals extend their wings fully forward as opposed to backward, forming an umbrella-like shape with both wings.

#### b. Physiological behaviours

This category characterizes behaviours occurring in response to changing environmental conditions (induced by external factors) or that are crucial to maintaining vital physiological processes (basal metabolism, internal core temperature).

1. **Ruffle**. Ruffling is indicated by a visible body-volume ratio increase (e.g. ruffled plumage), with extremities and bare parts drawn back or covered actively by the surrounding feathers. Performed habitually in response to low temperatures (average temperature below 10 °C). Activity levels are evidently decreased to prevent redundant metabolic energy loss and ensure a more efficient conservation of the body's core temperature. Could be an indicator for secondary pathology if the behaviour persists for a prolonged period and is not associated with external environmental changes.

2. **Heat-exposure display**: Opposite to ruffling; describes a reaction to high heat or sun exposure. The plumage is visibly slicked, and the body is extended vertically. While remaining perched, the carpal joints are held away from the body. In exceptional cases, the beak is widely open, with the respiratory rate evidently increased, indicated by a rapid horizontal movement of the tongue in quick succession (panting).

3. **Drink**: Describes the consumption of fluids. Drinking generally follows a bimodal pattern, with a peak in drinking intensity between 07:00 and 08:59, and another peak between 16:00 and 17:59. Drinking occurs more regularly during hot days or in response to excessive physical activity (up to five times/day) during the non-breeding season. Water intake often occurs before or after feeding, or soon after fresh water is provided. During water intake, the beak is positioned about 1 cm under the water surface, and water is carried out by alternating, rapid vertical movements of the tongue. Water is stored in the lower beak. After the lower beak is filled, the head is drawn slightly up, and the neck and forehead are flipped slowly backwards, so the water flows down the esophagus. These steps are usually repeated three to five times for sufficient water intake. Daily water intake or requirements were not measured in detail; however, observations made during the daily feeding schedules indicate that the approximate daily water requirements range between 8–12% of body mass, varying in response to changing nutritional conditions and abiotic factors.

4. **Food intake**: Two different types of food intake can be distinguished. Direct food intake is the method that birds usually use for food components with an average size <1 cm in diameter. For food components larger than ca. 1 cm in diameter, the food intake is facilitated by involving one leg (Figure S5c) to position and hold the food item. The second type of intake often involves multiple steps of manipulation or food preparation prior to the actual ingestion. The beak is essential for breaking the hard endocarp of nuts and for portioning a large food item into smaller fragments or processing parts of it. Both males and females showed similar lateral preference of a specific leg used to hold food items (i.e. footedness) (N_lefty_male_ = 21, N_righty_male_ = 13, N_lefty_female_ = 28, N_righty_female_ = 10, proportion test, Χ^2^ = 0.689, p = 0.407), with a clear bias towards using the left leg (exact binomial test, proportion of left leg = 0.681, CI_95%_ = 0.561–0.786, N_total_ = 72, *p* = 0.00294).

5. **Defecation**: This behaviour is occasionally performed intentionally in response to a mild disturbance and is then observed prior to a direct approach of an animal keeper or potential threat. It also occurs often during handling of neonates, but not during handling of adults, which usually defecate instead prior to or after handling. More commonly, defecation is an auto-maintenance behaviour. During defecation, the bird lowers its upper body and raises its tail up.

#### c.  Locomotion behaviours

Three different types of locomotion patterns were observed:

1. **Move**: While perched, individuals use a pattern of lateral leg displacement to achieve movement along horizontal perches, initiated by a movement of leg A, with leg B following a short interval and placed parallel to leg A (see Figure A6b). This pattern is repeated multiple times to ensure movement along horizontal elements. In contrast, a different pattern is used for directional movement on the ground, with legs moving forward in alternating order (i.e., as humans normally walk).

2. **Climb** (Figure A6a): Climbing involves multiple locomotor steps, where the beak is used predominantly as a third limb to lift or swing the body in the targeted direction. The mechanism of climbing on mesh is illustrated below; it is initiated by a disposition of the leg toward the target direction, followed by the beak, which helps push the body toward the same direction. The complete step is accomplished by the final disposition of the second leg (B), while the body is held stationary with the assistance of leg A and/or the beak. These steps are continued and repeated until the target location is reached. Modification of the above-described pattern is required to accommodate divergent spatial circumstances, e.g., climbing in dense vegetation, where a more complex approach is needed.

3. **Flying** (Figure A6c): The flight patterns of the Spix's Macaw are characterized as highly agile and capable of performing quick evasive maneuvers even in small spaces. Fledged chicks can acquire an extraordinary confidence in flight within a period of only 2–3 weeks of practice (post-fledging). For short flight distances, birds require a high wing-beat frequency to maintain consistent speed, while preferring to glide for longer distances (which minimizes energetic output), with the flight pattern interspersed with stronger wing beats. In the wild, birds perform multiple loops with strong but silent wingbeats before perching; turnovers are performed with high control over speed and directionality. Speed is adjusted just before landing. In the wild, flight patterns can vary, depending on the ethological context. Alert flights are usually characterized by erratic takeoffs, in which birds of the same social unit rapidly rearrange themselves in flight and follow the same direction, accompanied by alert calls. The end of an alert flight typically corresponds to the groups perching far from the take-off location, usually in a well-covered canopy of a tall tree. Foraging flights can be either directional or commuting, with linear or circling patterns, probably dependent on food abundance. While linear flight patterns may indicate that food resources are predictable and likely used at known feeding sites, circling flights allow for a larger search area, which should help locate potential food resources. Accordingly, circling flights are often observed in the dry season, probably in response to food shortages.

#### d. Inactivity behaviours

1. **Perch**: This behaviour does not involve any disposition or relocation of the body, with the bird remaining stationary using either one or both legs to perch.

2. **Resting** (Figure A5a): This behaviour represents the predominant activity pattern during the active daytime period, where individuals perch either partially isolated (inter-individual distance exceeds one body length) or near their mate or social partner (e.g., flocking birds) in a relaxed body posture. While resting, one leg is usually drawn back into the plumage.

3. **Roosting** (Figure A5b): Roosting can be subclassified into two different time-framed types: diurnal roosting and nocturnal roosting. Younger birds aggregate into flocks or groups if more than one bird is present in the aviary. Individuals in a communal flock leave and rearrange their perches in accordance with other members of the flock (centralized flock dynamics). However, individual activities during the diurnal period were non-directional and decentralized relative to the main flock.

Diurnal roosting is carried out in a more decentralized pattern by single individuals or small groups than nocturnal roosting, without necessarily linking these activities to flock members' actions. Our observations indicate that the species demonstrates high roosting site fidelity during the night, i.e., pairs or flocks were observed using the same perches as roosting sites for many consecutive weeks or months (if no changes were made by the keepers). The same behaviour was observed in the wild, where pairs or single birds would roost at the same sites, or even at the exact same location, for consecutive months. In contrast to nocturnal roosting sites, no site fidelity was observed during diurnal roosting. The duration and schedule of diurnal roosts vary between individuals and are context-dependent. Diurnal roosts observed without any notable external disturbance had an average duration of 20.75 ± 9.4 min (9.02–35.58; *n* = 13).

#### e. Agonistic behaviours

Intimidatory, aggressive, and conflict behaviours are summarized here under the term *agonistic behaviour;* descriptions are expanded from Marcuk *et al*., (2020).

1. **Neck and head feather raise**: The feathers of the head, neck, wings, and back are raised.

2. **Foot-lift**: Describes a slow, usually sideward directed presentation of one leg, with front toe pointed out towards the opponent, in a wave-like motion.

3. **Bill gape**: A wide opened bill presented towards the opponent or aggressor, usually with an evident mock bite display.

4. **Wing-raise display**: Show a bilateral wing unfolding either partly or fully with open bill, facing the opponent.

5. **Lunge**: Outline a thrust or lunge usually with closed bill towards opponents’ head, leg or upper body parts with a clearly visible mock bite display.

6. **Bite**: The aggressor directs the bill wide open toward the head, leg or upper body regions of the opponent or targets the nearest body point.

7. **Bill fence**: Reciprocal bill thrusting, where aggressor lunges or thrusts towards the opponent's head and opponent responds vice versa.

8. **Claw**: A physical socio-negative interaction involving the claw, where either the aggressor pushes one leg against opponents' upper body or reciprocal clawing occurs.

9. **Rush**: The aggressor walks rapidly, with wide open bill, in the direction of the opponent.

10. **Flying approach**: The aggressor lands directly on or a few centimeters away from the opponent.

11. **Flight attack**: The aggressor attacks subordinate in flight with a wide-open bill, and both claws usually directed towards the opponent's head.

12. **Fight:** A physical encounter with high intensity aggression involving two opponents with majority of the above-described forms of physical socio-negative interactions.

13. **Redirected aggression:** Redirection of mild to high intensity aggression from dominant partner to subordinate mate, when the potential intruder is unreachable. Can in occasional cases result from stereotypical behaviour, where one bird attacks the bird performing stereotypical behaviour.

#### f. Displacement behaviours

For time-activity patterns, these behaviours were categorized under submission or agonistic behaviour; descriptions are expanded from Marcuk *et al*., (2020).

1. **Displacement preen**: Performed as an exaggerated preening movement with a noticeable overexcitement in the form of alternating head shakes that intersperse the preening act.

2. **Displacement food-intake**: Forcefully executed, partial intake of a randomly chosen food item in front of an intruder.

3. **Displacement rub**: A pretended beak rubbing demonstrated as a part of a territorial display, performed in an exaggerated way, with beak rubbed on a solid surface in all available directions.

4. **Displacement scratch**: A rapid, alternating movement of the upper claws that in common form intends to remove foreign matter from head and neck region.

5. **Displacement hold-bite**: A redirected vigorous bite on a perch or other solid surface used for distraction during territorial encounters.

6. **Displacement head down shake**: A degraded or partly executed head shake, usually performed on one side.

7. **Displacement yawn**: Identical to the common form; an extension of upper and lower beak under maximal contraction of the mandibular muscles; with the head slightly withdrawn back.

8. **Displacement allopreening**: A pseudo socio-positive interaction where one individual starts to preen another individual during a territorial encounter.

9. **Displacement mutual feed**: A pseudo mutual feeding initiated without evident passing pre-digested food from donor to acceptor.

10. **Irritated body shake**: An exalted version of the ordinary body shake, execution accelerated and accomplished by combined eye blazing (i.e., pupil contractions) and jerking.

11. **Bill clasp**: short convergent interlock of the beaks of two individuals.

#### g. Submission behaviours (expanded from Marcuk et al., 2020):

1. **Turn away**: Aggression avoidance by turning away from aggressor without a body disposition.

2. **Slide away**: Describe the recede from an aggressor by moving or flying away with active body disposition.

3. **Alert and fear reaction**: The body axis arranged nearly vertically along the perch; the individual remains motionless with eyes wide open, and plumage sleeked.

4. **Apparent death display**: An auto-defense display used by chicks during the post-natal period as an anti-predatory strategy that feigns muscular rigidity and post-mortal motionless.

5. **Bob**: A cyclic repetition of head downward jerk, following a jerk in opposite direction.

6. **Head-tilt solidarity display**: The head moved to one side, slowly withdrawn and tilted backwards, following an oscillation to the other side in a waving motion.

7. **Crouch-quiver solidarity display**: The subordinate assumes a hunched posture accompanied by alternating wing-quivers, interspersed eventually with head shakes and submissive calls.

8. **Upside-down lift solidarity display**: Individual climbing on the roof of an aviary or in the canopy of a small tree or bush, lifting the body axis and hanging either with one leg or both legs on the mesh or on a twig. Legs are moved under the wing over the head.

9. **Peer**: A mutual convergent head downward jerk, with one head side directed towards the source of disturbance.

10. **Unison jerk**: A polyfunctional, highly synchronized display given in unison by a pair or social unit with an initial vertical extension of the body axis, habitually accomplished by a partly wing unfold and high-pitched call in unison.

11. **Singleton jerk**: identical to unison jerk, thus performed only by a single bird.

#### h. Social behaviour

The Spix's macaw is a gregarious species that engages in frequent social interactions with its partner or flock members. This set of behaviours encompasses all behaviours involving socio-positive interactions with a conspecific throughout the year, except for behaviours displayed only during the breeding season.

1. **Contact-sitting** (Fig. A7b): Describes the perching in very close proximity or direct contact with the social partner, observed often before or after allopreening sessions or during roosting.

2. **Mutual nibbling** (Fig. A7a): Outlines a form of mutual maintenance targeting the peripheral area of the beak, bare parts, and auricular patch, where usually both individuals simultaneously remove scales or dirt from the aforementioned areas.

3. **Allopreening** (Fig. A7d): Allopreening includes all forms of mutual or reciprocal preening involving two or sometimes also three individuals (flocks/parents preening a chick), where the social partner actively and carefully removes feather dust or any foreign material by gently placing the feathers in the beak, pulling the feather through the beak, and co-temporally using the tongue to clean the respective plumage. A few types of allopreening can be distinguished, like the (i.) head allopreening, where bird A usually targets to preen the head, neck, and nape of bird B, mostly remaining perched vertically, occasionally placing the facing leg on the back, without grabbing it. (ii.) Under wing allopreening; where bird A moves its head to gently lift one wing of bird B and to preen the underwing and abdominal plumage (iii.). Back allopreening describes preening of the dorsal plumage along the back towards the upper tail coverts.

4. **Reciprocal cloacal preen** (Fig. A7c): A typical form of allopreening, where both partners preen the undertail coverts and cloacal region, occasionally also the rectrices of the partner in aligning the body in contrasting directions.

5. **Non-sexual social play**: This form of social play was observed among all age groups except for old non-breeding adults. Social play encompasses a wide variety of behaviours that are usually performed in an erratic and exciting manner. These behaviours often include agonistic actions, but do not display any intention to injure or harm the counterpart. Social play can involve a single individual, in which case it is referred to as "social object play" (Diamond and Bond, 2003). In this case, there is no physical interaction with other group members. Social object play always involves interaction with the environment (e.g., a toy) and often results in group members mimicking the play or engaging in the second type of social play, which involves at least two individuals: "direct social play". This latter form of social play corresponds to a mutual, vigorous interaction involving repeated lunging, biting toes, and grabbing flight feathers, wings, legs, or tail feathers (i.e., "play fighting", as described by Diamond and Bond in 2003). Social play also includes foot-lifting, clawing, and other intimidating behaviours sometimes referred to as "play chasing" (Diamond and Bond, 2003). However, agonistic behaviours always serve a strictly non-aggressive purpose. Social play intensifies when accompanied by behaviours such as wing-raising, frantic movement along perches, and neck-twisting. Occasionally, individuals become entangled with their playing partner while hanging from the roof or perch. Social play can result in a loss of balance, causing one or both individuals to fall from the perch. Social play also occurs in parent-chick interactions. Although we only observed it during the post-fledging period, social play may also occur, unrecorded, inside the nest.

6. **Begging** (Fig A8a): Two forms of begging can be distinguished. One is observed in psittacine neonates, while the other is seen in adult females. Chicks begin exhibiting typical begging behaviour less than one hour after hatching. They emit a strong, high-pitched begging call that undergoes several changes during postnatal growth. Small chicks flick their wings powerfully and bob their heads while begging, but they cannot hold their heads upright for long. Fledged chicks remain perched while begging, showing a submissive body posture and softly bobbing their heads in the direction of one of their parents. They flick their wings but rarely extend them fully and habitually uplift their carpal joints while simultaneously emitting the begging call. Although begging triggers parental feeding, chicks tend to continue begging even after becoming independent or receiving sufficient food from their parents. Conversely, the second type of begging is observed in adult females and is directed toward their male partner, primarily during incubation and chick rearing, and occasionally during the pre-egg laying period to encourage mutual feeding. This behaviour is structurally similar to that of a fledged chick. We assume that it probably serves a sexual purpose during the early pre-egg laying period and a clear nutritional purpose during the later nestling stages. We also observed context-independent begging in adult Spix's macaws, which often indicated pathologies, such as weight loss or poor food intake, and required immediate intervention.

7. **Non-sexual allofeeding**: This behaviour of food transmission occurs between a mother and her chick, or between mates, but it typically remains unidirectional during a single feeding event. The only exception seems to occur during the first days after hatching; we observed parents performing reciprocal feedings before feeding the chick. On some occasions, such reciprocal exchanges occurred multiple times in a row, involving the transfer of predigested food. It is unclear whether this behaviour aims to provide proenzymes and probiotics from both parents to the chick to promote intestinal flora and immune system development. In general, three types of allofeeding were observed: Type I involves feeding, in which the male is the food donor feeding the female during different nestling stages. Once the female can feed herself, she sometimes feeds the male, who is also taking care of the chicks. Allofeeding type II occurs only as a parent-chick interaction, in which predigested food is passed from one parent to a chick. Allofeeding type III is less frequently observed. It occurs between siblings inside the nest or after fledging. Predigested food is passed from one sibling to another, either after food intake or after feeding. In this case, we cannot confirm whether food is successfully transferred or whether mutual feeding is simply a display, since the behaviour is very brief.

#### i. Sexual behaviours

1. **Sexual social play**: It is structurally very similar to the non-sexual social play, as it includes the same arrangement of agonistic behaviours; however, this type of social play is exclusively found in newly formed pairs, young pairs, or pairs during the onset of the breeding or pre-egg-laying period. The male and female often enter the nest together, either without an evident trigger or following a copulation or mutual feeding. Allopreening precedes sexual social play, which is initiated by either the male or female. The initiator lunges towards the partner in a waving motion or softly bites and pulls its toe, and the partner reciprocates. Sexual social play is often short (< 2 min) and intensifies rapidly. During this time, one or both partners may fall on their back and gently grab and claw at their partner's wings, head, or feet. Sometimes, the entanglement escalates into short sessions that resemble a fight. After the play fight, however, both partners suddenly become completely still, lying on their backs. After a few seconds, they turn around, either engaging in allopreening, or repeating the social play, or leaving the nest.

2. **Apparent copulation:** These differ from the pseudo-copulations of *Anodorhynchus* spp., and they do not represent incomplete copulations or attempts at copulation. They are more frequently observed in younger pairs or at the beginning of the breeding season, before the first copulations are initiated. As in true copulations, this behaviour is initiated by the female. The female aligns her body horizontally or diagonally while remaining perched. Unlike true copulation, this behaviour is more like "mutual forward leaning" in which both the male and female arrange their bodies parallel to each other while remaining perched for a short time, merging their cloacal regions without establishing true cloacal contact.

3. **Jerk display**: Similar to the unison jerk but performed mostly in unison. It most often happens following sexual allofeeding, but it can also occur during the pre- or post- copulatory context. Both males and females show synchronized eye blinking (i.e., pupil contractions) before performing the typical jerking pattern by lifting their body. This behaviour is usually performed silently, not necessarily accompanied by a call. Short in duration, averaging 3.11 ± 1.13 s (1.6–6.24, *n* = 18).

4. **Preening display**: It is very similar to displacement preening, which can sometimes be observed during the early stages of pair formation. Preening display most frequently occurs in the form of pre-copulatory behaviour and rarely follows copulation. It is performed in unison while both adults are perched in close proximity. The male and female begin to preen their back or belly erratically, accompanied by eye-blazing and excited calls.

5. **Scratch display:** This is a pre- or post-copulatory behaviour in which the male and female begin scratching their heads in unison or alone. Duration shorter than ordinary scratching: 2.69 ± 0.37 s (2.08–3.2, n = 11).

6. **Rapid turn**: The male and female turn in an exaggerated manner multiple times on a perch. It represents a typical pre-copulatory behaviour. During copulation, a similar behaviour can sometimes be observed where copulation is interrupted, and the male and female rearrange their positions along the perch by turning around before proceeding with a second attempt at copulation.

7. **Sexual allofeeding** (Fig. A8b): As for the non-sexual allofeeding, this behaviour involves the intentional passing of predigested food from the donor to the acceptor by interlocking the beaks of both individuals. The food donor regurgitates the food, which is passed through rapid horizontal tongue movements to the lower beak of the partner, while both beaks remain locked. Sometimes accompanied by head-bobbing. In sexual allofeeding, the two involved individuals are the two mates. In some Neotropical psittacines, the male is typically the food donor (Lantermann, 1987; Snyder et al., 1987), whereas for Spix's macaws, we observed reciprocal feeding, with both sexes acting as donors and receivers. We observed either the male or the female initiating the feeding or exchanging food, with roles often swapping across consecutive feeding bouts. Two types of sexual allofeeding were recorded: one that occurs during the pre-egg laying period and a second, prolonged type that occurs closer to the egg-laying period. The first type of sexual allofeeding rarely occurs soon before or after copulation. Detailed observations of a single pair showed that a sexual allofeeding consisted of 1–4 single feedings performed 5–22 s apart. Single feeding events within a feeding session lasted, on average, 8.25 ± 3.66 s (2.08–21.44, *n* = 64). We recorded an average of 4.61 ± 2.57 regurgitations (range 0–9, *n* = 49) before or during feeding bouts. No quantitative data was recorded for head bobbing, which was completely absent in some pairs. Single feeding events where the female fed the male were slightly longer, 8.73 ± 4.50 s (2.56–21.44; *n* = 34), than single feeding events from the male to the female: 7.71 ± 2.36 s (2.08–12.0; *n* = 30), but they do not differ significantly (Mann Whitney U test, z = 0.26, *p* = 0.795). A second type of sexual allofeeding, showing a prolonged duration averaging 76.16 ± 40.99 s (35.14–163.68, *n* = 8), was recorded closer to egg-laying for the same pair. The purpose of this behaviour remains unclear, as it is unlikely that food was passed on for such a long time period, even when starting with a full crop.

8. **Copulation** (Fig A9): Typical side-to-side copulation as in other macaws. The female usually initiates copulation following an allopreening session (Fig. A9a), which mostly targets the head and sometimes the underwing or belly. The female eagerly pushes her lower body towards the male, aligning her body diagonally or vertically with her tail facing up and her head facing down. The female signals her readiness (Fig. A9b) for copulation by emitting a repetitive soft cracking call and tilting and pushing her head toward the male repeatedly. Incomplete copulations are mostly terminated at this phase, during which the male remains stationary for 1–2 minutes without actively engaging in the process. In some cases, the male and female require several attempts to initiate copulation without completing it. The active phase is initiated by the male (Fig. A9c), who carefully establishes cloacal contact by moving the leg closest to the female under her wing and over her back. The foot does not grasp the back, but rather lies on top of it, with the toes contracting in slow pulses while copulation takes place. The male positions his tail and lower body diagonally, opposite to the female. He maintains his balance by moving his upper body from side to side, or by biting into the perch. Occasionally, he holds the position by fluttering with both wings (Fig. A9d) or interlocking the beak with the female (Fig. S9e). The male maintains cloacal contact with pulsing movements while the female remains stationary and only gently moves her lower body. Throughout the entire copulation process, the soft-cracking calls intensify until copulation is complete. The final phase is defined as the termination phase. First, the pulsing movements and calls stop. The male and female remain motionless in the copulation position for 5–15 s (Fig. A9f). Usually, after this time, the female dismounts and leaves the copulation site first (Fig. A9g), perching close to the male. Autopreening and shaking are often seen after a successful copulation. Incomplete copulations are short-duration and don't feature the three aforementioned phases. Complete copulations of 12 pairs that were recorded on video and analyzed had an average duration of 179.80 ± 60.98 s (70.33–352.04, *n* = 924). In contrast, incomplete copulations lasted on average 53.37 ± 22.36 s (19.04–106.24, *n* = 62) and were often followed by another copulation attempt.

#### j. Behavioural disorders

None of the "behavioural disorders" were observed in any of the birds after releasing were released into the wild.

*Non-physical stereotypic behaviour (displays)*:

1. **Erratic flights**: Erratic short flights, usually performed in loops, with birds repeatedly flying circles around the same spot, accompanied occasionally by short sessions with the crouch-quiver display. Chronic forms were not observed; mostly triggered by the presence of a keeper.

2. **Head tilt** (see category g., behaviour 6): Not observed in a chronic form; in the majority, triggered by the presence of a keeper, performed for a prolonged period until the potential intruder leaves the proximity of the aviary.

3. **Crouch-quiver solidarity display** (see g., behaviour 7): Prolonged and chronic forms (occurring without an obvious trigger and performed for long periods, occasionally accompanied by submission calls)

4. **Upside-down lift solidarity display** (see g., behaviour 8): Prolonged and chronic forms, frequently accompanied by the crouch-quiver solidarity display.

5. **Loop-walking**: Describing a repeated motion sequence, without flying, usually by walking in a circulating pattern, without an obvious direction or functional context.

*Physical behaviours* (involve intentional physical interaction):

6. **Pterotillomania (Feather-plucking behaviour)**: Defines a periodic or chronic form of feather damage behaviour. Sudden sequences of feather damage behaviour observed in adults or juveniles can be associated with sudden stress, interspecific aggression, post-breeding stress, or pathology. Usually, stress-related cases tend to bite off their primaries and secondaries, rather than pluck feathers. Plucking can be periodic and reversible, or non-reversible if performed over a longer period, with affected areas staying bare. Plucking of the plumage feathers in chicks by the females was recorded in most females at ACTP Germany (75%, 18 out of 24), while in none of the parent-raised females that reared chicks successfully.

7. **Overt allopreening**: Describing the intentional over-preening and plucking of feathers, mostly affecting the front head, neck or wing pin feathers of the current partner. Rarely chronic forms, often observed during breeding season or in the form of post-breeding stress.

8. **Auto-mutilation**: Describes any extraordinary, rare form of intentional infliction of wounds, documented only in three birds. Requiring immediate intervention, as self-mutilation often leads to life-threatening injuries or secondary infections.

9. **Allo-mutilation**: Very rare adverse behaviour, where one bird inflicts injuries to the mate by either aggressively removing feathers, scales, or skin fragments, without preceding socio-negative interactions (observed in four pairings).

10. **Redirected aggression**: While mate aggression is considerably rare, some males redirect aggression in the form of lunging, biting, clawing, or bill gaping towards the partner in the proximate presence of an animal keeper. Escalations into fights are extraordinarily rare.

11. **Egg destruction**: Observed only during the breeding season. However, observed in several females who intentionally destroyed individual eggs or the entire clutch shortly after laying or after several days. Not a native behaviour, often the result of induced stress (e.g., taking eggs, psychological trauma). Not observed in males, also not in young males, while that behaviour was observed in immature males of Lesser Antillean Amazons and Lear's Macaws paired with an older, egg-laying female. Sometimes the female stopped incubating for a couple of hours before destroying the eggs.

12. **Infanticide**: Very rare. Some females showed adverse behaviour by mutilating the chick's beak, wing tips, and toes, often leading to a lethal outcome for the chick within the first 12 hours after hatching. Instantaneous killing of chicks is extremely rare (documented in three females at ACTP Germany and in three other females in Al Wabra Wildlife Preservation = AWWP and appears to be associated with extrinsic stress).

### (3) Comments

Our study provides the first description of the full suite of behaviour, including behavioural disorders, for the Spix's macaw.

The qualitative descriptions for the maintenance, locomotion, resting, and physiological behaviours provided here for the Spix's macaw are consistent with behaviours reported for other macaws, for species with both smaller and larger body sizes (Uribe, 1982; Christiansen and Pitter, 1992; Pitter and Christiansen, 1995, 1997; Barros, 2001; Schneider *et al*., 2006; Favoretto *et al*., 2024).

Descriptions of the activity, maintenance, and locomotion behaviour for other Psittaciformes suggest, in general, inter-generic consistency in functional context and motoric execution. A possible explanation is that these behaviours evolved in ancestors to perform a specific arrangement of elementary tasks—without undergoing much change in their functionality over subsequent generations. These include plumage maintenance, stretching, or head scratching. Slight dissimilarities observed in such basal behaviours, as in the over-the-wing head-scratch in *Nestor* spp. and under-the-wing head-scratch in *Agapornis* spp. (Dilger, 1960; Higgins, 1999), could arise from adaptations to distinct morphological (body size, shape) and anatomical characteristics differing between species, without changing the ethological context of the specific behaviour.

Climbing and general movement (Uribe, 1982; Rowley, 1990; Higgins, 1999) or flight patterns are well documented for neotropical parrots (Whitney, 1996) and do not differ from the description provided here. The locomotion of psittacines, in general, differs from that of many other avian families because of the zygodactyl arrangement of the toes and the use of the beak as an accessory element in maneuvering their motoric steps (Blanco *et al.,* 2018). This specific arrangement evolved independently in other extant avian families, such as Cuculidae, Galbulidae, Bucconidae and the Pici (Johansson and Ericson, 2003). The footedness and preference for using one specific leg have previously been discussed in detail for parrots (Harris, 1989) and also occur in Spix's macaws. Submission, displacement, and antagonistic behaviours will not be discussed in more detail, as they were reviewed in detail by Marcuk *et al.,* (2020).

Bathing behaviour as recorded here has been qualitatively described for only a few psittacines in captivity, with no data available for most "macaws" regarding latency to bath, refractory period, and duration of baths. Baths lasted on average 10.8 ± 1.7 min in a study on the Orange-winged Amazon (*A. amazonica*; Murphy *et al.,* 2011), which is within the reported range we recorded for the Spix's macaw. The other elements of bathing behaviour mentioned here were also observed in closely related species (*Ara* spp., *Primolius* spp., *Orthopsittaca manilata*; VM. *pers. obs*.).

Social play has been documented in several psittacines (Diamond and Bond, 2003), while sexual social play has not yet been documented for macaws. Social play between siblings, parents, and chicks or young birds in a flock is probably linked to different stages of social development (e.g., neophilia and sociable behaviour), whereas sexual social play seems to contribute more to intra-pair harmony and is probably a vital element of the pair formation process. As reported above, this behaviour was observed exclusively during the pre-egg-laying period, mostly hidden inside the nest. That would also probably explain the lack of records for wild individuals. The authors (pers. obs.) have also recorded sexual social play in other macaws in human care, including *Ara ambiguus, A. chloropterus, A. glaucogularis, A. macao, A. araurana, A. militaris mexicanus,* and *Primolius maracana.*

We also documented here for the first time sexual displays that superficially resemble displacement behaviours, observed mostly in response to mild or severe disturbance or conflicting situations. Displacement behaviours and displays were also documented in pre-copulatory or post-copulatory contexts in lorikeets (*Trichoglossus* spp., Serpell, 1979), whereas none were described in macaws.

Copulations of most neotropical parrots, particularly macaws, have only been qualitatively described for a few species. In general, copulation seems to be structurally consistent (e.g., side-to-side copulation) between closely related taxa. Our description aligns with that provided for *Ara macao cyanoptera* (Inigo-Elias, 1996)*, A. militaris mexicanus* (Reyes Macedo, 2007)*, A. araurana* (Bianchi, 1998)*, Primolius maracana* (Barros, 2001), *Anodorhynchus* *leari* (Favoretto et al., 2024)/*hyacinthinus* (Schneider et al., 2006). We also observed similar patterns for *A. glaucogularis,* *A. macao macao*, *A. chloropterus, A. severa, A. rubrogenys, A. ambiguus, Primolius couloni* and *Diopsittaca nobilis nobilis*. According to the literature, durations range from 5–15 seconds for *Ara macao* (Inigo-Elias, 1996)*,* 30–40 seconds (in the wild, Bianchi, 1998) or 30 seconds (in captivity, Locatelli et al., 2013) for *A. ararauna,* while 'courtship behaviour', like mutual feeding and copulations has been reported to last up to 15 minutes combined for *A. glaucogularis* and *A. rubrogenys* (Robiller, 1999; Abramson *et al*., 1995). These records differ from those we recorded for Spix's macaws, as well as for Hyacinth and Lear's macaws (see Marcuk *et al*., 2025). Conversely, detailed records for the Mexican military macaw (*Ara militaris militaris*) seem to resemble what we observed for Spix's macaws, with one study stating the copulation duration range as 2–5 min (*n* = 4, Carreón, 1997) and another with more accurate data as 1.75 ± 1.30 min (0.03–4.30, *n* = 28, Reyes Macedo, 2007). The very short durations probably also included durations of incomplete copulations. Captive *Primolius auricollis* were recorded to copulate up to 7 min (Robiller, 1999), and copulation durations of ca. 6 min were also recorded for the Spix's macaws. These comparisons should be interpreted with caution since available data on copulation durations seem to be based on different, usually not explicit definitions of the copulation event. In particular, it is often unclear whether the values given refer to the duration of a full copulation or only part of it (both can greatly differ if a copulation is interrupted). Some publications may have reflected the active copulation phase (cloacal contact), while others may have included the duration of pre-copulatory displays as well. Therefore, we strongly encourage authors to be explicit when recording quantitative data on complex behaviour to establish meaningful reference values for different species.

### (4) Additional references

1. Abramson, J., Speer, B.L., Thomsen, J.B., 1995. The large macaws: Their care, breeding and conservation. Raintree Publications.
2. Barros, Y.M., 2001. Biologia comportamental de Propyrrhura maracana (Aves, Psittacidae): Fundamentos para conservação in situ de Cyanopsitta spixii (Aves, Psittacidae) na caatinga (Doctoral dissertation). Universidade Estadual Paulista, Brazil.
3. Bianchi, C., 1998. Biologia reprodutiva da arara-canindé (Ara ararauna, Psittacidae) no Parque Nacional das Emas, Goiás (Master's thesis). Brazil.
4. Blanco, G., Hiraldo, F., Tella, J.L., 2018. Ecological functions of parrots: An integrative perspective from plant life cycle to ecosystem functioning. Emu - Austral Ornithol. 118(1), 36–49. https://doi.org/10.1080/01584197.2017.1387031
5. Carreón, A.G., 1997. Estimación poblacional, biología reproductiva y ecología de la nidificación de la guacamaya verde(Ara militaris) en una selva estacional del oeste del estado de Jalisco (Bachelor's thesis). Universidad Nacional Autónoma de México, Mexico City, Mexico.
6. Christiansen, M.B., Pitter, E., 1992. Aspects of behaviour of juvenile red-fronted macaws (Ara rubrogenys) in the wild. Le Gerfaut 82, 63–69.
7. Diamond, J., Bond, A.B., 2003. A comparative analysis of social play in birds. Behaviour 140, 1091–1115. https://doi.org/10.1163/156853903322589650
8. Dilger, W.C., 1960. The comparative ethology of the African parrot genus Agapornis. Z. Tierpsychol. 17, 649–685. https://doi.org/10.1111/j.1439-0310.1960.tb00412.x
9. Favoretto, G.R., Sugieda, A.M., Guida, F.J., Piratelli, A.J., 2024. Ethogram of Anodorhynchus leari (Aves, Psittacidae) in captivity: A framework for behavioral studies. Iheringia. Sér. Zool. 114, e261579. https://doi.org/10.1590/1678-4766e261579
10. Harris, L.J., 1989. Footedness in parrots: Three centuries of research, theory, and mere surmise. Can. J. Psychol. 43(3), 369–396. https://doi.org/10.1037/h0084225
11. Higgins, P.J. (Ed.), 1999. Handbook of Australian, New Zealand & Antarctic birds. Vol. 4: Parrots to Dollarbird. Oxford University Press.
12. Inigo-Elias, E.E., 1996. Ecology and breeding biology of the scarlet macaw (Ara macao) in the Usumacinta drainage basin of Mexico and Guatemala (Doctoral dissertation). University of Florida, USA.
13. Johansson, U.S., Ericson, P.G., 2003. Molecular support for a sister-group relationship between Pici and Galbulae (Piciformes sensu Wetmore 1960). J. Avian Biol. 34(2), 185–197. https://doi.org/10.1034/j.1600-048X.2003.03103.x
14. Lantermann, W., 1987. Die Blaustirnamazone – Biologie, Ethologie, Haltung. Urania-Verlag.
15. Locatelli, A.C., Wrublack, S.C., Basile, L.F., do Nascimento, A.F., Berber, G.D.C.M., de Araujo Berber, R.C., 2013. Comportamento reprodutivo e materno de araras-canindé (Ara ararauna) mantidas em cativeiro para conservação. Comunicata Sci. 4(4), 316–323.
16. Marcuk, V., Purchase, C., de Boer, D., Bürkle, M., Scholtyssek, K., 2020. Qualitative description of the submission and agonistic behavior of the Spix's macaw (Cyanopsitta spixii, Wagler 1832), with special reference to displacement displays. J. Ethol.38(2), 253–270. https://doi.org/10.1007/s10164-020-00650-6
17. Marcuk, V., Drigon, C., Gupta, B.K., Patil, S., Vercillo, U., Purchase, C., et al., 2025. First photographic records of aberrant white feathers in the threatened Lear's macaw Anodorhynchus leari, with further notes on its breeding behavior. Braz. J. Biol. 85, e296852. https://doi.org/10.1590/1519-6984.296852
18. Murphy, S.M., Braun, J.V., Millam, J.R., 2011. Bathing behavior of captive orange-winged Amazon parrots (Amazona amazonica). Appl. Anim. Behav. Sci. 132(3–4), 200–210. https://doi.org/10.1016/j.applanim.2011.04.010
19. Pitter, E., Christiansen, M.B., 1995. Ecology, status and conservation of the red-fronted macaw Ara rubrogenys. Bird Conserv. Int. 5(1), 61–78. https://doi.org/10.1017/S0959270900002951
20. Pitter, E., Christiansen, M.B., 1997. Behavior of individuals and social interactions of the red-fronted macaw Ara rubrogenys in the wild during the midday rest. Ornitol. Neotrop. 8(2), 133–143.
21. Reyes Macedo, G., 2007. Biología reproductiva de la guacamaya verde (Ara militaris) en la Cañada Oaxaqueña, Reserva de la Biosfera Tehuacán–Cuicatlán. Mexico.
22. Robiller, F., 1990. Papageien: Handbuch der Vogelpflege. Vol. 3: Mittel- und Südamerika. Deutscher Landwirtschaftsverlag.
23. Rowley, I., 1990. Behavioural ecology of the galah (Eolophus roseicapillus) in the wheatbelt of Western Australia. Surrey Beatty & Sons.
24. Schneider, L., Serbena, A.L., Guedes, N.M.R., 2006. Behavioral categories of hyacinth macaws (Anodorhynchus hyacinthinus) during the reproductive period at South Pantanal, Brazil. Rev. Etol. 8(2), 71–80.
25. Serpell, A.J., 1979. Comparative ethology and evolution of communicatory behaviour in the lorikeet genus Trichoglossus (Doctoral dissertation). University of Liverpool, UK.
26. Whitney, B.M., 1996. Flight behaviour and other field characteristics of the genera of neotropical parrots. Cotinga 5, 32–42.
27. Uribe, F., 1982. Quantitative ethogram of Ara ararauna and Ara macao (Aves, Psittacidae) in captivity. Biol. Behav. 7, 309–323.

**Figures**

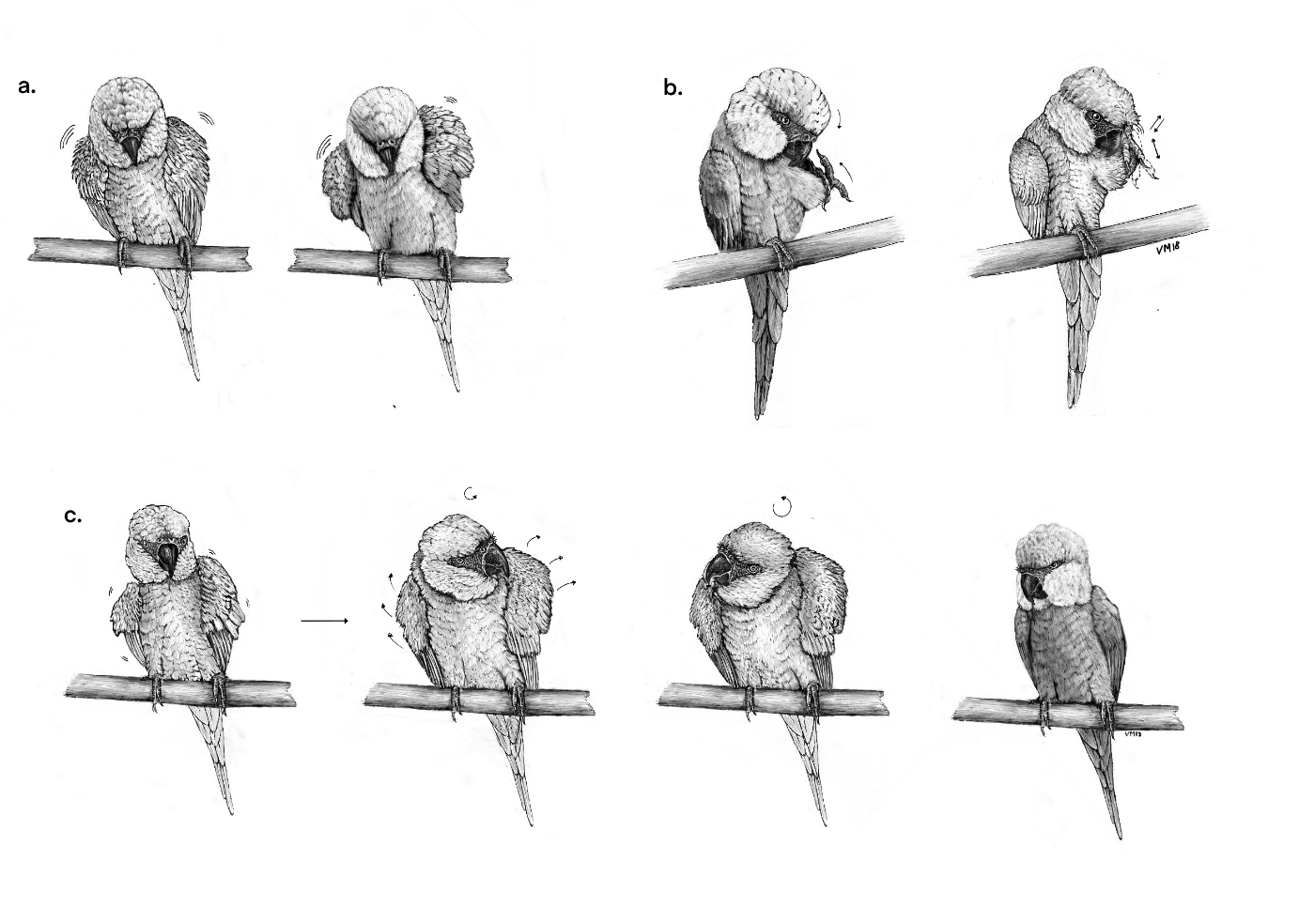

**Figure A1** – (a) body-shake (b) scratch (c) head-shake (all illustrations by corresponding author).

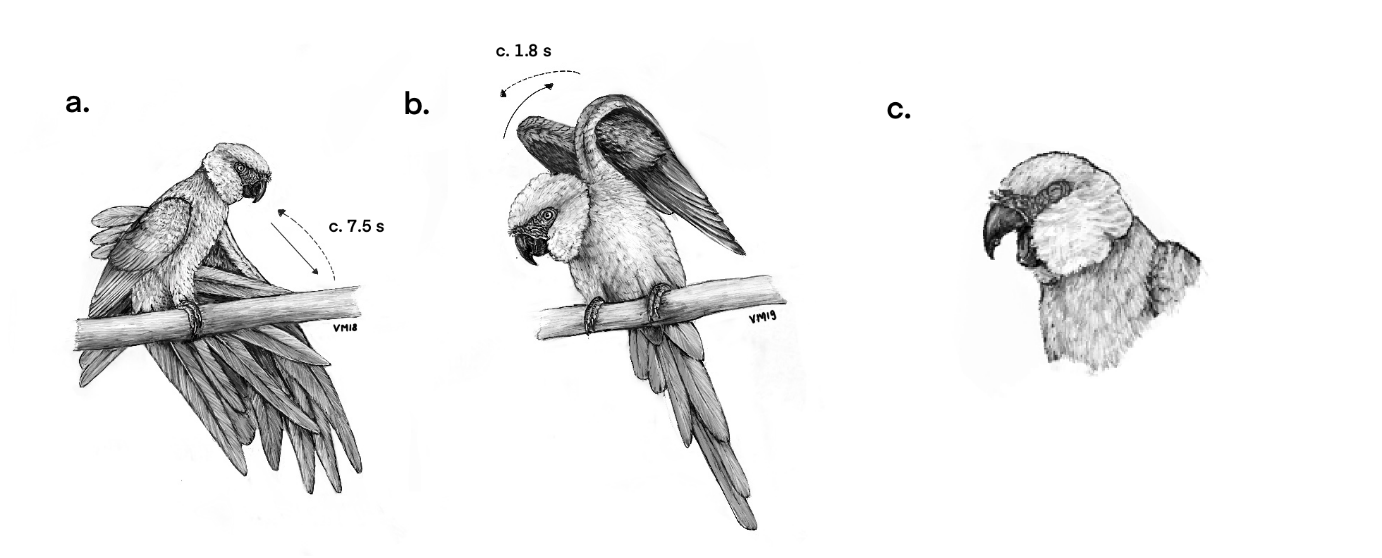

**Figure A2** – (a) wing & leg stretch (b) bilateral wing stretch (c) yawn.

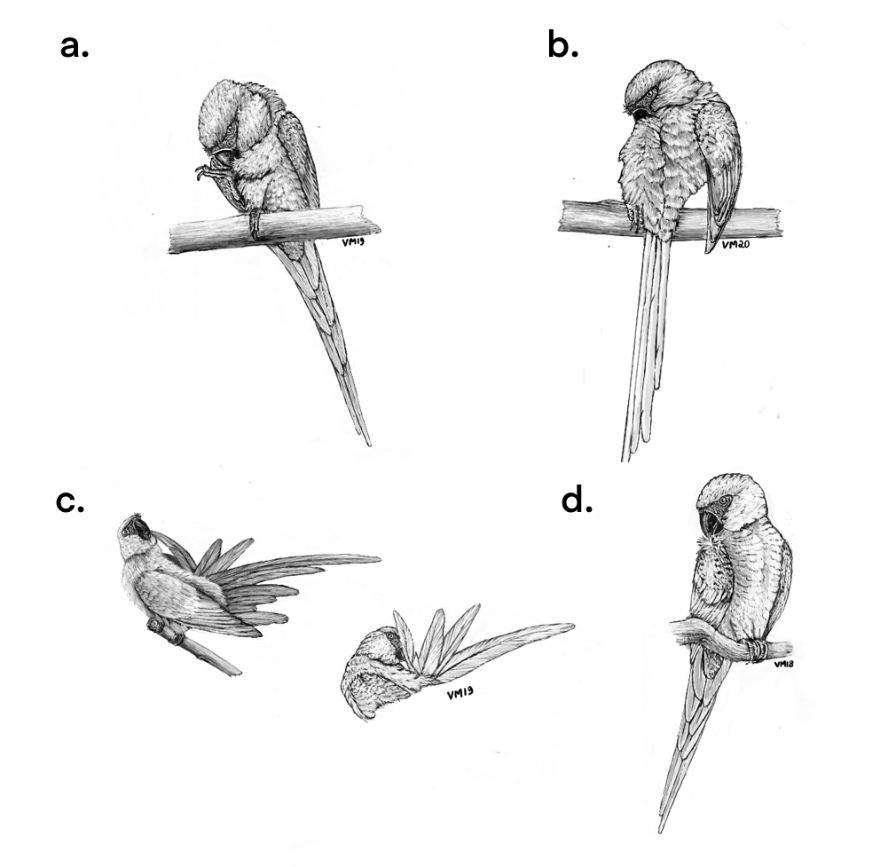

**Figure A3** – Distinct behaviours of auto-preening (a) touch-foot (b) back-preen (c) tail-preen (d) wing-preen.

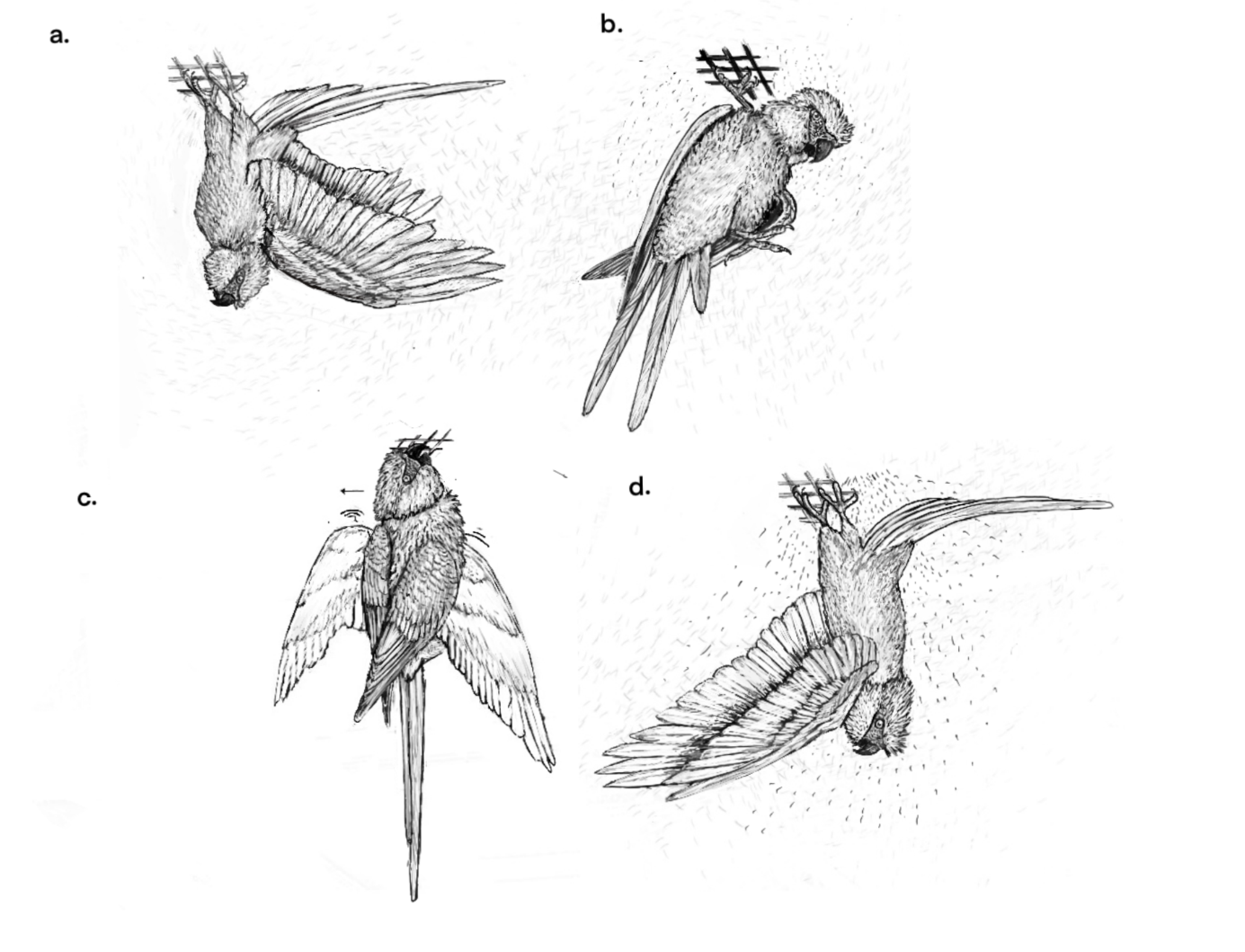

**Figure A4** – (a) backside hang (b) upside hang (c) beak hang (d) umbrella posture.

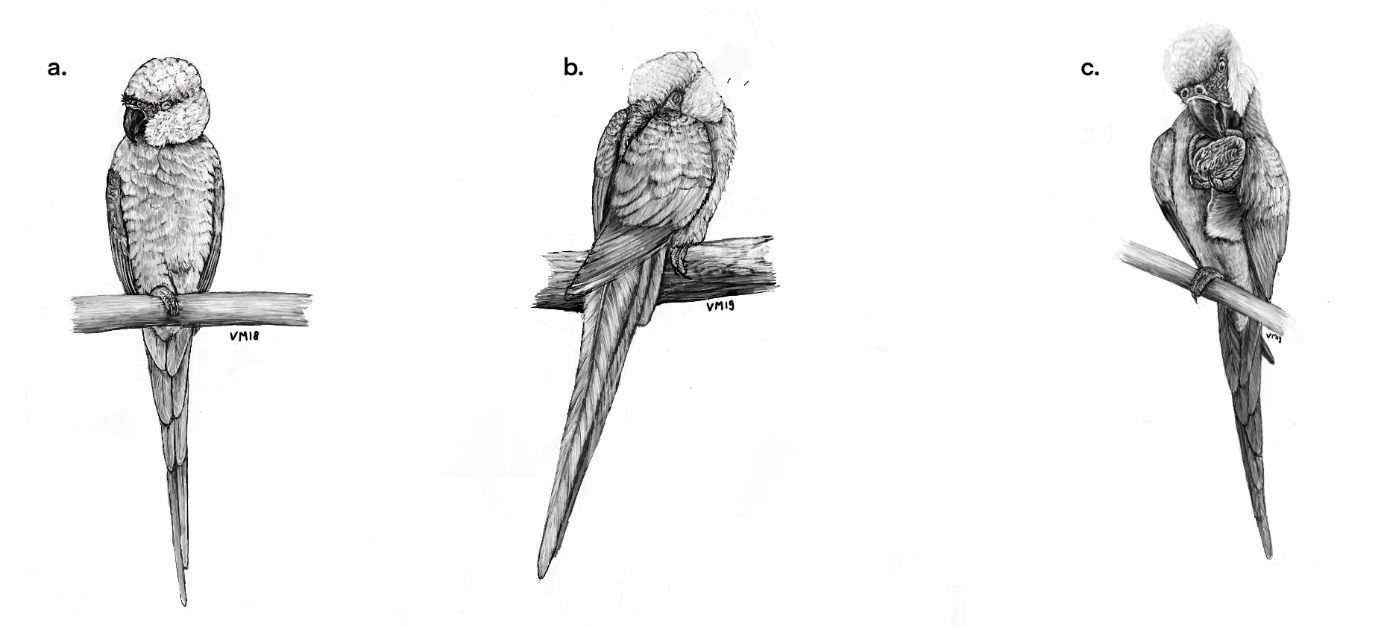

**Figure A5** – (a) rest (b) roost (c) typical facilitated foot intake supported by one leg**.**

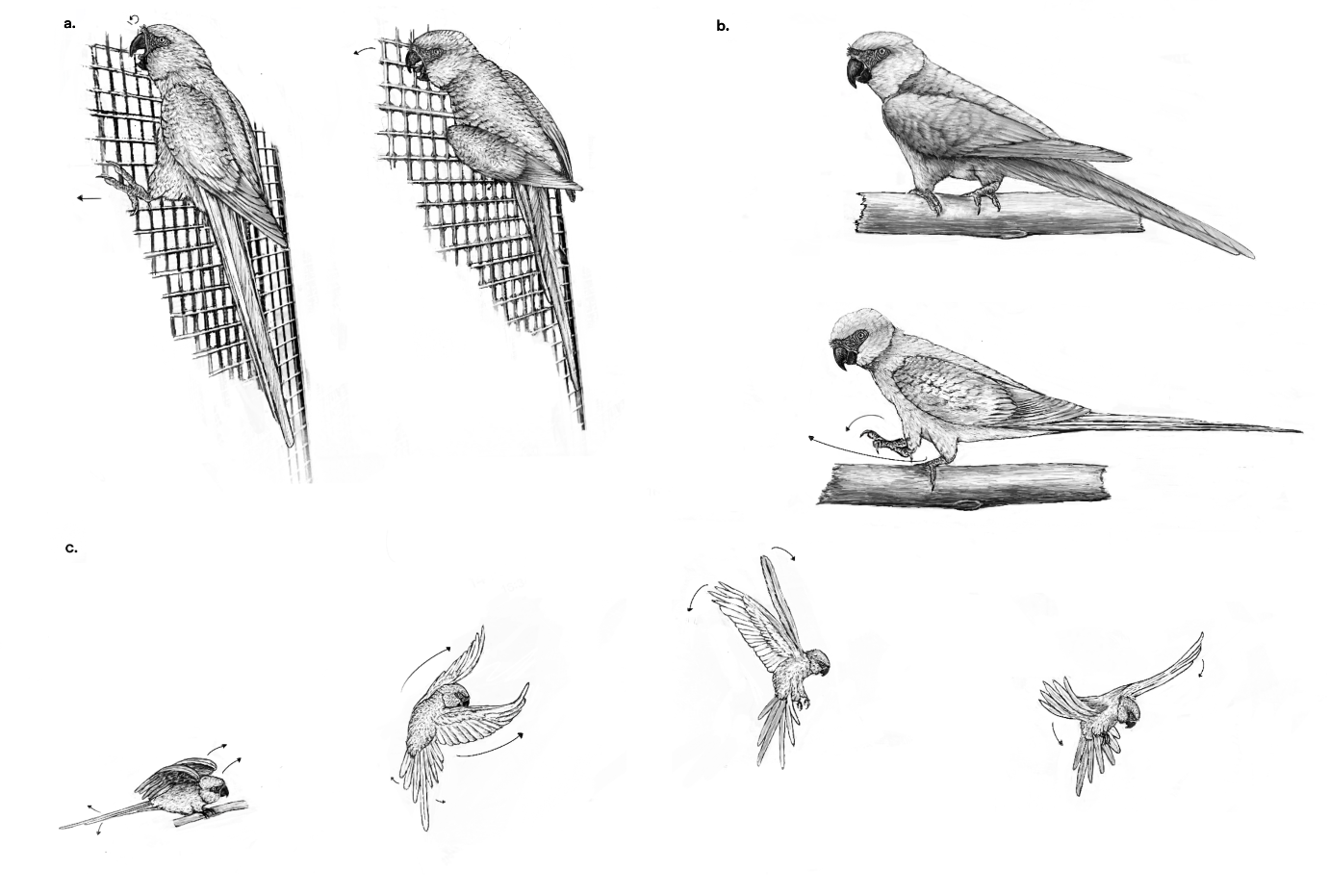

**Figure A6** – (a) climb (b) move (c) flight.

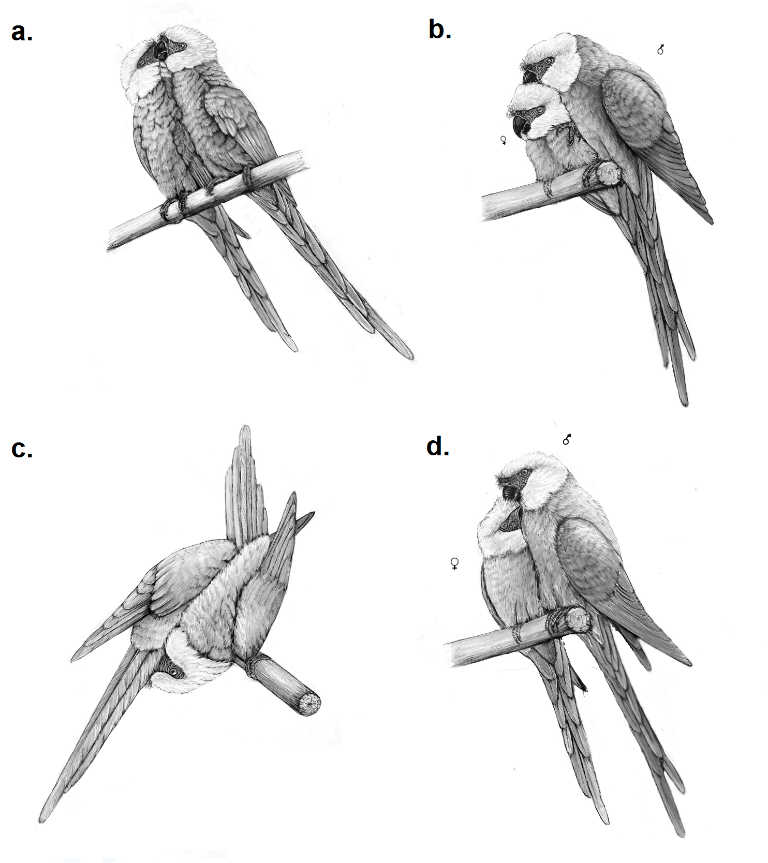

**Figure A7 –** (a) nibbling (b) contact-sitting (c) reciprocal cloacal preening (d) allopreening.

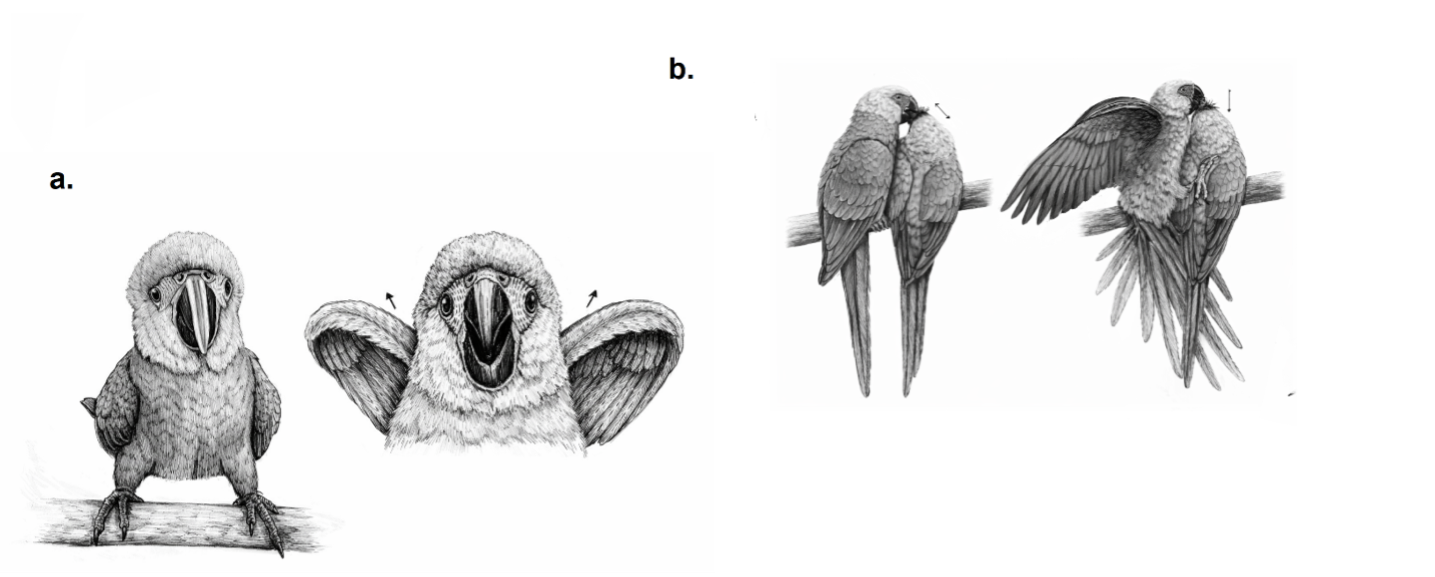

**Figure A8 –** (a) begging of a fledgling (b) sexual allofeeding.

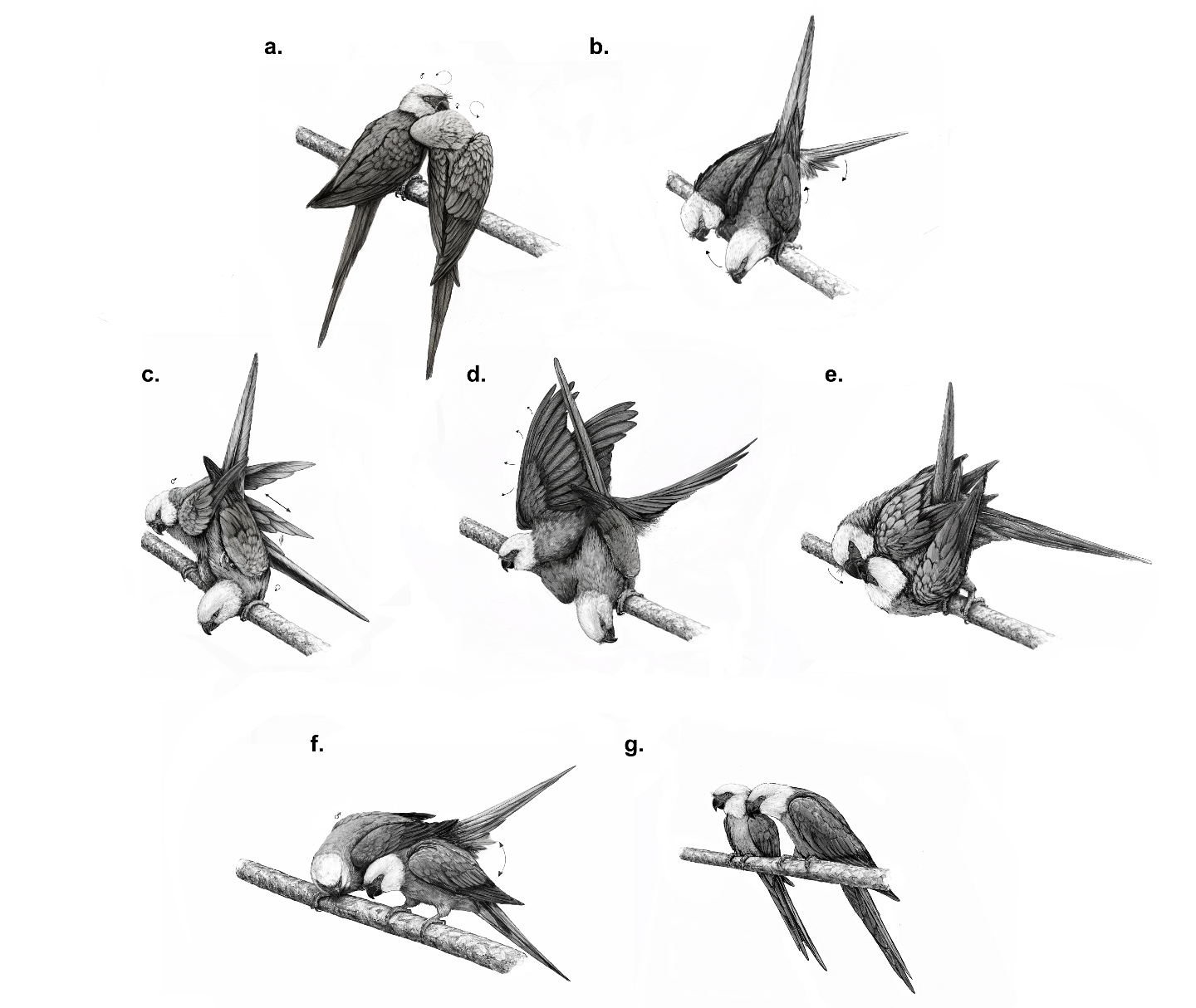

**Figure A9 –** Chronology of a typical copulation. Copulation initiation is illustrated in (a) and (b). The active copulation phase is given in (c), (d), and (e). Copulation termination is given in (f) and (g).

**Appendix II – Bootstrap analysis**

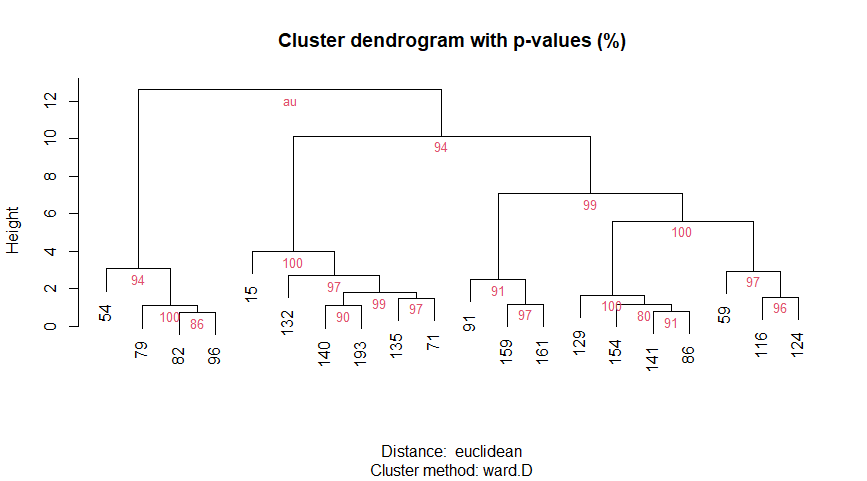

**Figure A10** – Bootstrap analysis was applied to the Ward-D clustering method. The dendrogram includes the studbook numbers associated with each individual and branch support. Generally, sub-clusters are well supported, while some higher-level clusters show moderate support, with AU values (percentage support of approximately unbiased tests) between 80 and 94%.
